# Systematic Screening of a Glycolysis-targeting Small-molecule Library Identifies Novel Inhibitors of Fungal Morphogenesis in *Candida albicans*

**DOI:** 10.64898/2026.04.23.720332

**Authors:** Dhrumi Shah, Yogita Martoliya, Sudharsan Mathivathanan, Ashish Shrivastava, Shailendra Asthana, Sriram Varahan

**Affiliations:** CSIR-Centre for Cellular and Molecular Biology, Hyderabad-500007, India; Academy of Scientific and Innovative Research (AcSIR), Ghaziabad-201002, India; BRIC- Translational Health Science and Technology Institute (THSTI), Faridabad-121001, India

**Keywords:** Fungal Morphogenesis, *Candida albicans*, Glycolysis, Pyruvate Kinase, Antifungal Drugs, Biofilm Formation, Murine Candidiasis

## Abstract

Fungal infections are an increasing global health concern, with *Candida albicans* emerging as a leading cause of mucosal and life-threatening systemic infections. *C. albicans* relies on yeast-to-hyphae transition, to establish systemic infections in the host. Our recent work demonstrated that glycolysis is a key regulator of hyphal differentiation in *C. albicans*. Leveraging this knowledge, we screened a small-molecule compound library containing glycolysis inhibitors for their ability to block fungal morphogenesis and identified multiple inhibitors of glycolysis that robustly block hyphal differentiation without compromising overall growth. While early glycolysis pathway inhibitors showed moderate effects, two compounds (NPD10084 and PKM2-IN-6) targeting pyruvate kinase activity emerged as the most potent inhibitors of fungal morphogenesis, completely blocking hyphal differentiation, in multiple filamentation-inducing conditions. Comparative RNA-Seq analysis revealed that pyruvate kinase inhibition using PKM2-IN-6 resulted in the repression of multiple genes associated with fungal morphogenesis, biofilm formation, and virulence. These transcriptional changes were accompanied by a significant reduction in biofilm formation and increased sensitivity to conventional antifungal drugs, including amphotericin B and fluconazole, in the presence of PKM2-IN-6. Furthermore, administration of PKM2-IN-6 significantly improved the survival of mice in a systemic model of murine candidiasis underscoring the in vivo efficacy of this compound. Molecular docking analysis showed stable binding of both compounds to the ATP-binding pocket of pyruvate kinase, suggesting ATP-competitive inhibition as the mode of action. Collectively, our study has identified novel inhibitors of fungal morphogenesis in *C. albicans* that target pyruvate kinase activity to increase antifungal susceptibility and attenuate the virulence of *C. albicans*.

**Importance:** *Candida albicans* is a major fungal pathogen that causes fatal infections, particularly in immunocompromised individuals. The increasing prevalence of antifungal resistance and the limited availability of effective antifungal therapies underscore the need for new treatment strategies. In this study, we identify pyruvate kinase, a key glycolytic enzyme, as an important regulator of fungal morphogenesis and virulence. Pharmacological inhibition of pyruvate kinase blocked hyphal differentiation without affecting fungal growth, impaired biofilm formation, and enhanced the activity of clinically relevant antifungal drugs. Transcriptomic analysis further revealed broad repression of genes involved in filamentation, biofilm formation, cell adhesion, antifungal resistance, and virulence, providing a molecular basis for these phenotypes. Importantly, pyruvate kinase inhibition also improved host survival in a murine model of systemic candidiasis. Together, our findings identify pyruvate kinase as a promising therapeutic target and demonstrate the potential of targeting fungal metabolism to effectively combat *C. albicans* infections.

## Introduction

Fungal infections represent a significant and growing public health concern, causing an estimated 1.7 million deaths annually and contributing to over 150 million severe cases worldwide (1–3). *Candida albicans* is one of the most common human fungal pathogens, causing infections collectively known as candidiasis that range from superficial to systemic infections. Mucosal infections, such as oropharyngeal and vulvovaginal candidiasis, are the most prevalent form of candidiasis (4, 5). In severe cases, *C. albicans* can cause life-threatening invasive disease, often originating from the gastrointestinal tract, damaged skin (e.g., burns), catheter-associated biofilms, surgical sites, or obstructed urinary tracts (6, 7). Reflecting its global health impact, the World Health Organization (WHO) has recently classified *C. albicans* as a critical priority human fungal pathogen (8, 9). Despite the substantial clinical burden caused due to *C. albicans* infections, effective treatment remains highly challenging. Many antifungal drugs exhibit significant toxicity to host cells, and the rapid emergence of drug resistance further compromises their efficacy (10). Consequently, current therapeutic options are limited and often insufficient, even in well-equipped healthcare settings (11). Collectively, these challenges emphasize the urgent need to identify new antifungal targets to support the development of safer, more effective therapeutic strategies. In this context, targeting fungal virulence mechanisms has gained considerable attention as an alternative therapeutic approach (12–15). One of the central virulence determinants of *C. albicans* is its ability to undergo morphological transitions from its yeast to hyphal form. Morphogenetic switching enables this pathogen to adapt to diverse host environments and facilitates multiple stages of infection, including tissue invasion, immune evasion, and biofilm formation (16, 17). Because these pathogenic processes are fundamentally dependent on filamentous growth, disrupting morphogenesis can directly impair the ability of fungi to establish and sustain infection. Importantly, targeting morphogenesis offers a promising anti-virulence strategy that may significantly reduce pathogenicity without necessarily affecting fungal viability, thereby potentially minimizing host toxicity and development of drug resistance associated with the use of conventional antifungals. Consequently, inhibiting fungal morphogenesis represents a highly attractive and strategically advantageous approach to combat recalcitrant fungal infections.

Given that transcriptional and signaling networks governing morphogenesis in *C. albicans* are well characterized, multiple groups have performed phenotypic screens and identified inhibitors of diverse signaling pathways, including Ras signaling, calcium homeostasis, and signaling pathways involved in the transcriptional regulation of morphogenesis, stress response, and virulence-associated gene expression (18, 19) that block hyphal differentiation in *C. albicans* (20, 21). Supporting this notion, a high-throughput screening approach identified filastatin, a small-molecule that inhibits adhesion, morphogenesis, and pathogenicity in *C. albicans* (22). Increasing evidence demonstrates that fungal morphogenesis in *C. albicans* is tightly integrated with central carbon metabolism. The transition from yeast to hyphal form requires extensive metabolic reprogramming to meet the elevated energetic and biosynthetic demands associated with polarized growth, cell wall remodeling, and invasive development (23, 24). Our recent work has demonstrated that glycolysis-dependent sulfur metabolism orchestrates fungal morphogenesis in *C. albicans* and this in turn is critical for its virulence (25). Consequently, this work adds to the growing literature that central carbon metabolic pathways like glycolysis, the tricarboxylic acid (TCA) cycle, and alternative carbon utilization pathways, play essential roles not only in energy production but acts as regulatory hubs that link nutrient availability to morphogenetic developmental programs (26–28). However, a systematic chemical evaluation of compounds that inhibit central carbon metabolic pathways including glycolysis and TCA cycle, and ultimately block morphogenetic switching, remains largely unexplored.

In this study, we performed a comprehensive functional screening of a commercially available glycolysis compound library to identify novel inhibitors of fungal morphogenesis that attenuate hyphal differentiation without compromising the overall growth of C*. albicans*. Notably, our results demonstrate that pharmacological inhibition of pyruvate kinase activity by NPD10084 (29) and PKM2-IN-6 (30) potently blocks hyphal differentiation of *C. albicans* across diverse filamentation-inducing conditions without affecting overall fungal growth, revealing a morphogenesis-specific metabolic vulnerability. The inability of exogenous pyruvate supplementation to rescue filamentation in the presence of these inhibitors further suggests that disruption of glycolytic flux, rather than pyruvate depletion alone, underlies this observed effect. To gain further mechanistic insights underlying these phenotypes, we performed comparative RNA-Seq analysis following pharmacological inhibition of pyruvate kinase using PKM2-IN-6, during hyphal differentiation. Transcriptomic profiling revealed extensive transcriptional remodeling, including coordinated repression of gene networks associated with filamentation, biofilm formation, cell adhesion, antifungal resistance, and fungal virulence, providing a mechanistic basis for how perturbation of pyruvate kinase activity attenuates hyphal differentiation in *C. albicans*. Consistent with our transcriptomic analysis, both the pyruvate kinase inhibitors significantly reduced biofilm formation and enhanced the antifungal activity conventional antifungal agents, including fluconazole and amphotericin B. Furthermore, administration of PKM2-IN-6 significantly improved survival of the mice in a systemic model of murine candidiasis, underscoring the in vivo efficacy of this compound. Overall, these findings establish selective inhibition of glycolytic activity as a promising antifungal strategy and identifies pyruvate kinase as a previously unexplored therapeutic target for attenuating morphogenesis-mediated fungal virulence.

## Results

### Screening and Identification of Glycolytic Inhibitors that Block Hyphal Differentiation in *Candida albicans*

In order to identify small-molecule inhibitors of glycolysis that specifically block fungal morphogenesis, we screened 112 compounds from the ‘Glycolysis Compound Library’ (MedChem Express HY-L058) that specifically interfere with distinct steps of glycolysis and glycolysis-related pathways (Fig 1A and 1B) (S1 Table). These compounds were categorized based on the glycolytic step or enzyme they target (Fig 1B). We first focused on compounds targeting glucose transporters (GLUTs), which mediate glucose uptake across the plasma membrane and constitute the primary entry point for glucose into the glycolytic pathway (31, 32). Accordingly, 23 GLUT-targeting compounds were screened for their ability to block hyphal differentiation in *C. albicans* wild-type (SC5314), under standard filamentation-inducing conditions (Spider medium). While none of these compounds caused a noticeable growth defect at the tested concentration of 250 µM, 12 compounds resulted in a marginal reduction in hyphal differentiation compared to the untreated control, suggesting that perturbation of glucose transport influences morphogenetic switching in *C. albicans* without compromising the overall growth (Fig 1C). We next focused on compounds targeting hexokinase/glucokinase, enzymes that catalyze the first committed and regulatory step of glycolysis by phosphorylating glucose to glucose-6-phosphate (33, 34). In total, 23 hexokinase/glucokinase-targeting compounds were screened, of which 6 moderately reduced the ability of *C. albicans* to undergo hyphal differentiation without compromising the overall growth with the exception of one compound (Globalagliatin), which also resulted in a growth defect (Fig 1D). As a positive control, we included 2-deoxy-D-glucose (2DG), a well-characterized glycolytic inhibitor that we have previously demonstrated to be a potent inhibitor of fungal morphogenesis in *C. albicans* (25). To assess whether inhibition of intermediate glycolytic steps impacts fungal morphogenesis, we next screened individual compounds targeting phosphofructokinase (PFK), glyceraldehyde-3-phosphate dehydrogenase (GAPDH), and phosphoglycerate kinase (PGK), for their ability to block hyphal differentiation. Interestingly, none of these inhibitors resulted in a significant reduction in hyphal differentiation compared to the untreated control (Fig 1E-1G). We then focused on enolase, a key glycolytic enzyme that catalyzes the conversion of 2-phosphoglycerate to phosphoenolpyruvate (PEP), a high-energy intermediate that directly feeds into the final step of glycolysis (33). A total of 5 enolase-targeting compounds were screened for their ability to block hyphal differentiation, of which one resulted in a modest reduction in hyphal differentiation relative to the untreated control (Fig 1H).

**Fig 1.**
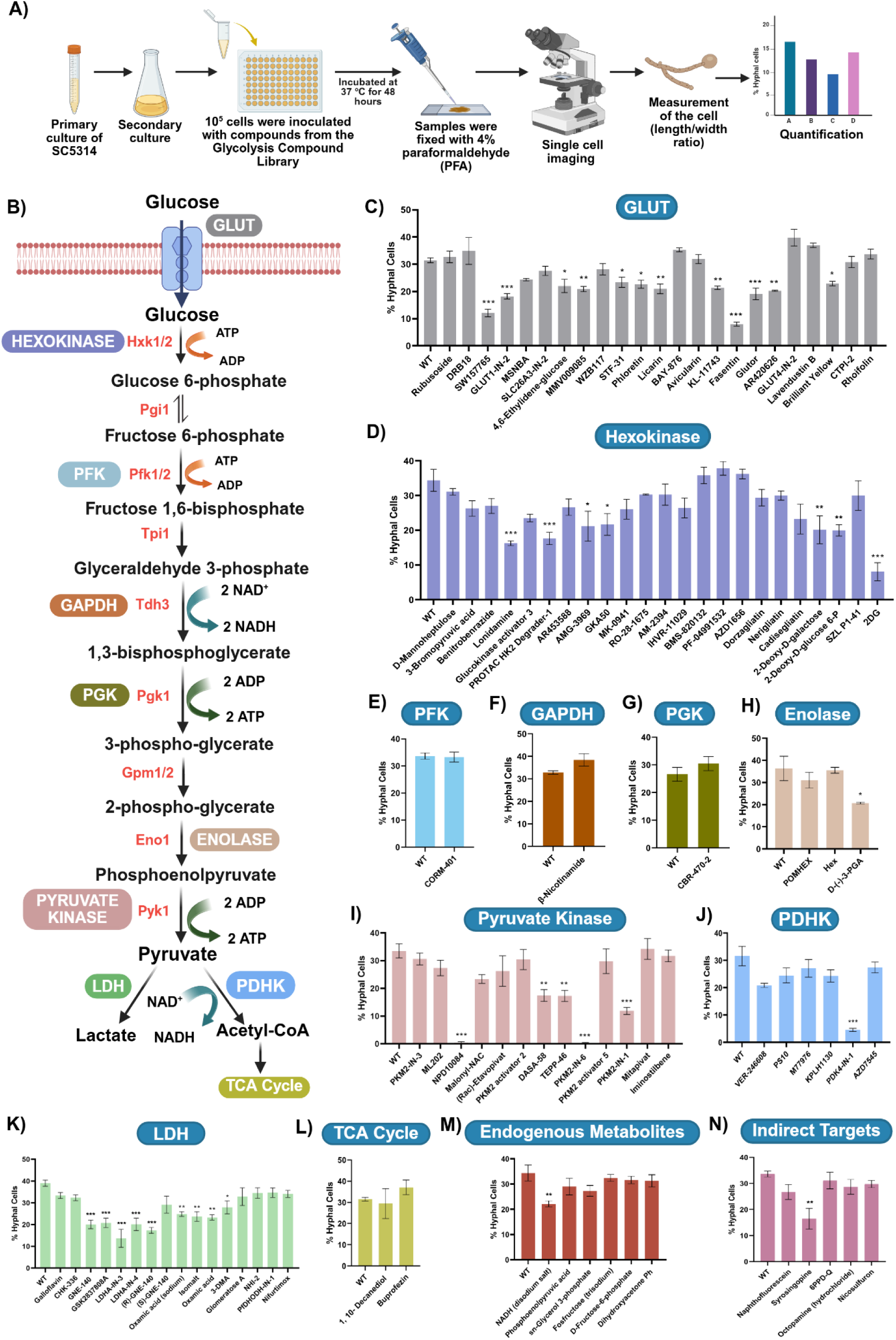
Screening and identification of glycolytic inhibitors that block hyphal differentiation in *C. albicans*. **(A)** Schematic overview of the experimental procedure for the filamentation assay conducted in a 96-well plate. **(B)** Schematic overview of glycolysis, highlighting the specific steps targeted by the inhibitors. **(C-N)** Quantitative estimation of hyphal cells in the wild-type strain was performed in the presence of glycolysis inhibitors at a concentration of 250 µM. A total of 10^5^ wild-type cells were cultured in Spider medium, with or without glycolysis inhibitors, at 37 °C for 48 hours. Following incubation, cells were fixed and imaged using a Zeiss Apotome microscope. The length-to-width ratio of individual cells was measured using ImageJ software, and the percentage of hyphal cells within the total population was determined. Cells exhibiting a length-to-width ratio of 4.5 or greater were classified as hyphal cells. For each inhibitor, more than 300 cells were counted. Statistical analysis was performed using one-way ANOVA, ***(P < 0.001), **(P < 0.01), and *(P < 0.1). Error bars represent SEM. **(C)** Quantitative estimation of hyphal cells in the presence of GLUT-targeting compounds at concentration of 250 µM. **(D)** Quantitative estimation of hyphal cells in the presence of hexokinase-targeting compounds at concentration of 250 µM. **(E)** Quantitative estimation of hyphal cells in the presence of PFK1 inhibitor at concentration of 250 µM. **(F)** Quantitative estimation of hyphal cells in the presence of GAPDH inhibitor at concentration of 250 µM. **(G)** Quantitative estimation of hyphal cells in the presence of PGK1 inhibitor at concentration of 250 µM. **(H)** Quantitative estimation of hyphal cells in the presence of enolase-targeting compounds at concentration of 250 µM. **(I)** Quantitative estimation of hyphal cells in the presence of pyruvate kinase-targeting compounds at concentration of 250 µM. **(J)** Quantitative estimation of hyphal cells in the presence of PDHK-targeting compounds at concentration of 250 µM. **(K)** Quantitative estimation of hyphal cells in the presence of LDH-targeting compounds at concentration of 250 µM. **(L)** Quantitative estimation of hyphal cells in the presence of TCA cycle-targeting inhibitors at concentration of 250 µM. **(M)** Quantitative estimation of hyphal cells in the presence of endogenous metabolites at concentration of 250 µM. **(N)** Quantitative estimation of hyphal cells in the presence of indirect targets of glycolysis at concentration of 250 µM.

Next, we screened compounds targeting pyruvate kinase, the terminal enzyme of glycolysis that catalyzes the conversion of PEP to pyruvate (35), for their ability to block hyphal differentiation. A total of 16 pyruvate kinase-targeting compounds were screened, of which 8 compounds significantly reduced hyphal differentiation and 3 of these compounds resulted in a growth defect. Notably, two compounds, NPD10084 and PKM2-IN-6, completely abolished hyphal differentiation at the tested concentration of 250 µM without compromising the overall growth of *C. albicans* (Fig 1I). These findings demonstrate that inhibition of pyruvate kinase activity potently suppresses hyphal morphogenesis in *C. albicans*, highlighting late glycolytic enzymes as promising targets for attenuating morphogenesis-mediated fungal virulence. Following our systematic screening of compounds targeting glycolytic enzymes, we extended our screening to central carbon metabolic pathways that link glycolysis to mitochondrial respiration, specifically focusing on pathways regulating acetyl-CoA production and its entry into the TCA cycle (36). To this end, we screened a total of 7 compounds targeting pyruvate dehydrogenase kinase (PDHK), a key regulatory enzyme that controls the activity of the pyruvate dehydrogenase complex thereby enabling the conversion of pyruvate into acetyl-CoA (37, 38). Among the 7 PDHK-targeting compounds screened, only one showed a significant reduction in hyphal differentiation compared to the untreated control (Fig 1J). In parallel, we also screened compounds targeting lactate dehydrogenase (LDH), an enzyme central to anaerobic metabolism that catalyzes the interconversion of pyruvate and lactate (39, 40), for their ability to block hyphal differentiation. A total of 20 LDH-targeting compounds were screened. Notably, 9 of these compounds caused a moderate reduction in hyphal differentiation compared to the untreated control, and 4 of these compounds resulted in a growth defect (Fig 1K).

In addition to compounds that directly target glycolytic enzymes, we also screened a subset of small-molecules that target other glycolysis-related metabolic pathways, such as the TCA cycle and endogenous metabolites of central carbon metabolism, for their ability to block fungal morphogenesis in *C. albicans*. These compounds are known to affect the expression of glycolytic enzymes, thereby indirectly impacting glycolytic flux (41–43). We screened a total of 2 TCA cycle-targeting compounds and 6 endogenous metabolites and out of those 8 compounds, only one compound resulted in a moderate reduction in hyphal differentiation compared to the untreated control (Fig 1L and 1M). Finally, we screened 7 compounds that indirectly target glycolysis for their ability to block hyphal differentiation. Out of these, 2 compounds showed a growth defect, and only one compound resulted in a moderate reduction in hyphal differentiation compared to the untreated control (Fig 1N). Furthermore, 13 compounds that significantly compromised the overall growth of *C. albicans* at 250 µM concentration were further evaluated for their ability to block filamentation at lower concentrations (100 µM and 10 µM). Out of the 13 compounds, 7 compounds significantly compromised overall growth even at 100 µM, and the remaining 6 compounds moderately reduced hyphal differentiation (S1A Fig). The 7 compounds that inhibited overall growth of *C. albicans* even at 100 µM, were further tested for their ability to block hyphal differentiation at a concentration of 10 µM. At this concentration, although none of the compounds compromised the overall growth of *C. albicans*, they did not affect hyphal differentiation (S1B Fig).

Collectively, our comprehensive screening identified 36 small-molecule inhibitors that reduced hyphal differentiation at 250 µM concentration, and 6 small-molecule inhibitors that significantly reduced hyphal differentiation at 100 µM concentration. However, only two compounds, NPD10084 and PKM2-IN-6, completely blocked hyphal differentiation without compromising the overall growth of *C. albicans*, (Fig 1I). Notably, both of these compounds belong to the pyruvate kinase–targeting group, underscoring the critical importance of this terminal glycolytic step in regulating morphogenetic switching in *C. albicans*. Based on their strong hyphal inhibitory activity, these two compounds were selected for further characterization.

### Dose-dependent Effect of Pyruvate Kinase Inhibitors on *C. albicans* Hyphal Differentiation

Based on our previous results, we also evaluated the dose-dependent effects of the two selected pyruvate kinase inhibitors, NPD10084 and PKM2-IN-6, by assessing their impact on hyphal differentiation at lower concentrations (100 µM and 50 µM). NPD10084 and PKM2-IN-6 are inhibitors of the human pyruvate kinase isoform PKM2 and target both its metabolic and non-glycolytic functions (29, 30). NPD10084 disrupts PKM2 interactions with β-catenin and STAT3, thereby attenuating downstream signaling (29), whereas PKM2-IN-6 (compound 7d), an imidazopyridine-based thiazole derivative, induces G2-phase cell cycle arrest and apoptosis, demonstrating significant anticancer potential, particularly in triple-negative breast cancer (30). We observed that NPD10084-treated *C. albicans* exhibited a significant reduction in hyphal differentiation at 100 µM, whereas this inhibitory effect was not observed at 50 µM. PKM2-IN-6 demonstrated a strong inhibitory effect on hyphal differentiation at both tested concentrations. At 100 µM, PKM2-IN-6 almost completely abolished hyphal differentiation. Notably, even at a lower concentration of 50 µM, PKM2-IN-6 significantly inhibited hyphal differentiation compared to the untreated control (Fig 2A). These results suggest that PKM2-IN-6 is a more potent inhibitor of *C. albicans* morphogenesis compared to NPD10084.

**Fig 2.**
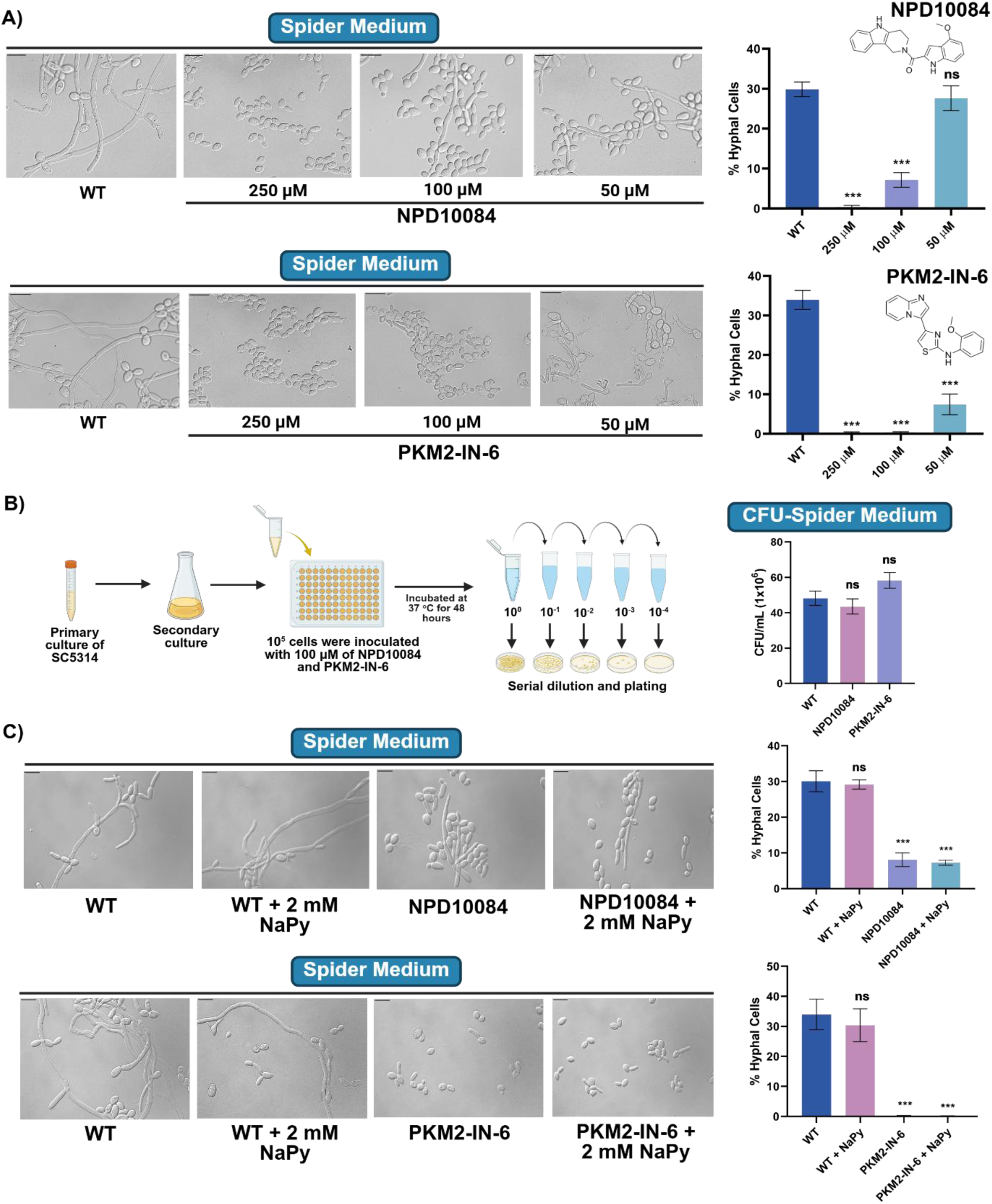
Dose-dependent effects of pyruvate kinase inhibitors on *C. albicans* hyphal differentiation and exogenous supplementation of downstream metabolite pyruvate fails to rescue this phenotype. **(A)** Quantitative estimation of hyphal cells in the wild-type strain was performed in the presence and absence of NPD10084 or PKM2-IN-6 at concentrations of 250 µM, 100 µM, and 50 µM. A total of 10^5^ wild-type cells were cultured in Spider medium with or without NPD10084 or PKM2-IN-6 at the specified concentrations, incubated at 37 °C for 48 hours. Cells were subsequently fixed and imaged using a Zeiss Apotome microscope. The length-to-width ratio of individual cells was measured using ImageJ, and the percentage of hyphal cells within the total population was determined. Cells with a length-to-width ratio of 4.5 or greater were classified as hyphal cells. For each condition, more than 300 cells were counted. Statistical analysis was conducted using one-way ANOVA ***(P < 0.001) and ns (non-significant). Error bars represent SEM. Scale bar represents 10 µm. **(B)** Schematic overview of the experimental procedure for the CFU assay, including CFU counts of *C. albicans* in the presence of 100 µM NPD10084 or PKM2-IN-6. Statistical analysis was performed using one-way ANOVA, ns (non-significant). Error bars represent SEM. **(C)** Sodium pyruvate (NaPy) add-back assay was conducted in the presence and absence of NPD10084 and PKM2-IN-6. Wild-type cells (10^5^ cells per well) were cultured in Spider medium supplemented with 2 mM NaPy and treated with either NPD10084 or PKM2-IN-6 at a final concentration of 100 µM. Plates were incubated at 37 °C for 48 hours to induce filamentation. After incubation, cells were fixed and imaged using a Zeiss Apotome microscope. The length-to-width ratio of individual cells was measured using ImageJ, and the percentage of hyphal cells in the total population was determined. Cells with a length-to-width ratio of 4.5 or greater were classified as hyphal cells. More than 300 cells were counted per inhibitor condition. Statistical analysis was performed using one-way ANOVA, ***(P<0.001) and ns (non-significant). Error bars represent SEM. Scale bar represents 10 µm.

To confirm that the observed inhibition of hyphal differentiation by these compounds at 100 µM concentration was independent of effects on overall fungal growth, we performed colony-forming units (CFU) assays using the wild-type strain in the presence of 100 µM NPD10084 and PKM2-IN-6, while the untreated wild-type strain was used as a control. CFU analysis revealed no significant difference in *C. albicans* growth between the inhibitor-treated and untreated controls (Fig 2B). These results indicate that, at the tested concentrations, both NPD10084 and PKM2-IN-6 do not impair fungal viability, and these effects appear to be specific to *C. albicans* hyphal differentiation.

### Exogenous Supplementation of the Downstream Metabolite Pyruvate does not Rescue the Filamentation Defects Caused by the Pyruvate Kinase Inhibitors

The pyruvate kinase enzyme catalyzes the final step of glycolysis, converting PEP and ADP to pyruvate and ATP (35). The enzyme plays a crucial role in controlling the overall glycolytic flux and ATP production (44). As pyruvate is a very dynamic metabolite that feeds into multiple metabolic pathways (45), we wanted to investigate whether the lack of pyruvate is the reason for the hyphal differentiation defect caused by the addition of NPD10084 and PKM2-IN-6. Therefore, we aimed to determine whether exogenous pyruvate supplementation could rescue this filamentation defect in *C. albicans*. In order to do this, wild-type *C. albicans* was grown in Spider medium, in the presence and absence of 2 mM sodium pyruvate (NaPy) along with the respective inhibitors. We observed that the exogenous supplementation of NaPy was unable to rescue the hyphal differentiation defect caused by the presence of NPD10084 and PKM2-IN-6 in *C. albicans* (Fig 2C). This suggests that hyphal differentiation defect cause by the perturbation of pyruvate kinase activity could likely be due to reduced overall glycolysis flux since it is known that the accumulation of PEP can competitively inhibit steps of phosphorylation and isomerization of glucose to fructose 6-phosphate and non-competitively inhibits phosphofructokinase and aldolase, which in turn can alter overall glycolytic flux (46, 47). PEP can also directly inhibit the upstream glycolytic enzyme triosephosphate isomerase (TPI) competitively, thereby suppressing glycolytic flux (48).

### NPD10084 and PKM2-IN-6 Block Hyphal Differentiation in Multiple Filamentation-Inducing Conditions

Standardized in vitro morphogenesis-inducing conditions, including RPMI, serum-containing medium, Lee’s medium, Spider medium, low glucose medium and N-acetylglucosamine–supplemented medium, provide diverse environmental and nutritional cues that activate distinct signaling pathways and regulatory networks required for hyphal differentiation in *C. albicans* (19, 49–54). Consequently, *C. albicans* exhibits variable morphological phenotypes across different liquid and solid growth conditions, reflecting the context-dependent regulation of the yeast to hyphal transition (55). To determine whether hyphal differentiation is inhibited by NPD10084 and PKM2-IN-6 across these different morphogenesis-inducing media, *C. albicans* wild-type was allowed to undergo filamentation in Lee’s medium, RPMI medium, YPD medium with 10% serum (YP + 10% serum), and SC medium with 0.1% glucose (SC + 0.1% glucose), in the presence and absence of these inhibitors. In Lee’s medium, the wild-type strain exhibited ∼70% hyphal differentiation, whereas treatment with NPD10084 reduced the proportion of hyphal cells to ∼25%. On the other hand, treatment with PKM2-IN-6 reduced the proportion of hyphal cells to ∼2% (Fig 3A). In RPMI medium, wild-type strain exhibited ∼81% hyphal differentiation, whereas treatment with NPD10084 reduced the proportion of hyphal cells to ∼28%, and treatment with PKM2-IN-6 reduced the proportion of hyphal cells to ∼4% (Fig 3B). Similar phenotypes were observed in SC + 0.1% glucose and YP + 10% serum, wherein the presence of NPD10084 and PKM2-IN-6 completely blocked hyphal differentiation in SC + 0.1% glucose (Fig 3C), whereas in YP + 10% serum, treatment with NPD10084 reduced the proportion of hyphal cells to ∼14%, and treatment with PKM2-IN-6 reduced the proportion of hyphal cells to ∼0% (Fig 3D). These results indicate that the inhibition of hyphal differentiation by these compounds is not media-specific, demonstrating their robustness at inhibiting fungal morphogenesis triggered by diverse environmental cues.

**Fig 3.**
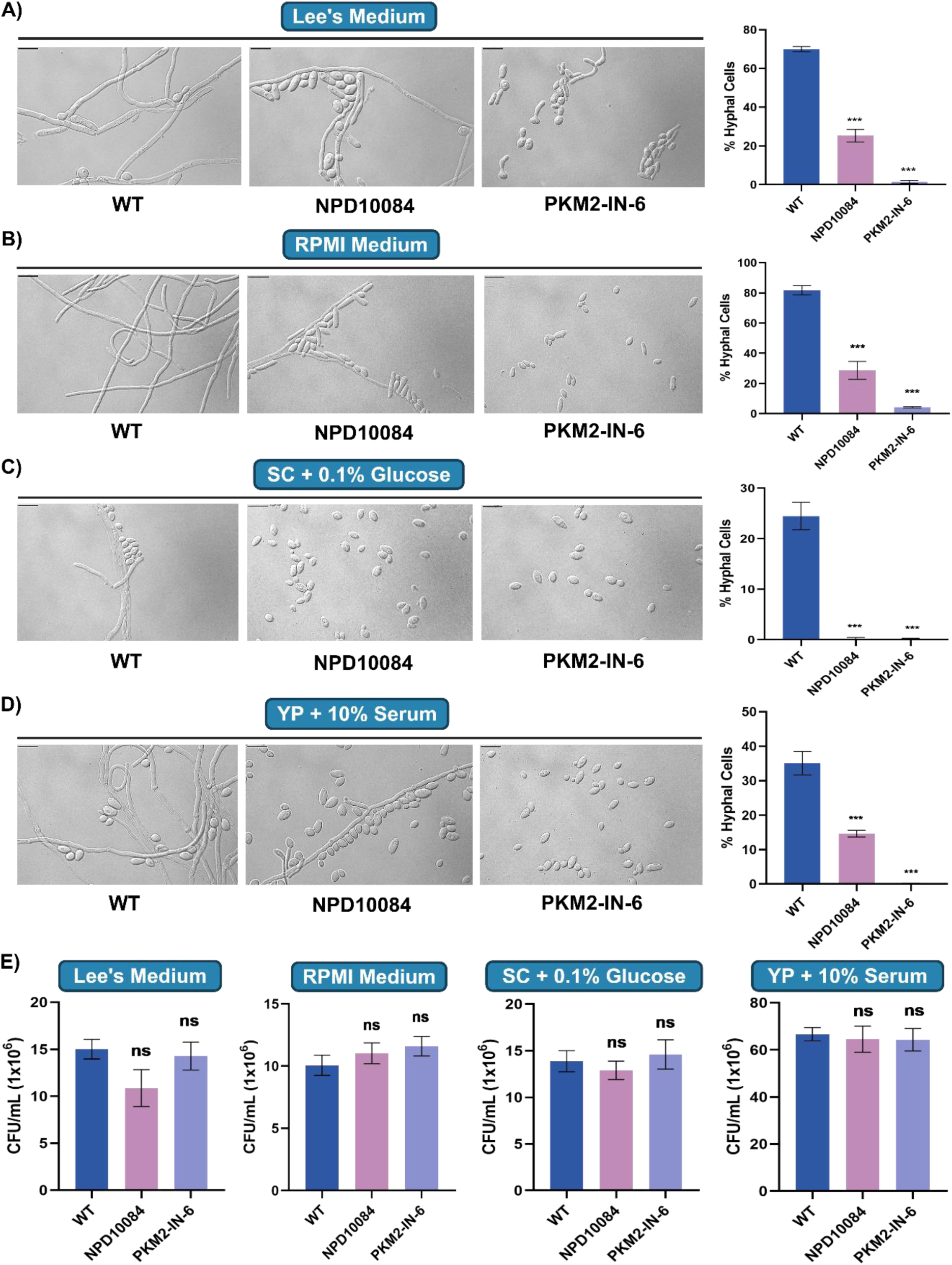
Pyruvate kinase inhibitors - NPD10084 and PKM2-IN-6 block hyphal differentiation in multiple filamentation-inducing conditions in *C. albicans*. **(A)** Quantitative estimation of hyphal cells was performed in Lee’s medium in the presence and absence of NPD10084 or PKM2-IN-6 at a concentration of 100 µM. Statistical analysis was conducted using one-way ANOVA, ***(P<0.001). Error bars represent SEM. Scale bar represents 10 µm. **(B)** Quantitative estimation of hyphal cells was performed in RPMI medium in the presence and absence of NPD10084 or PKM2-IN-6 at a concentration of 100 µM. Statistical analysis was conducted using one-way ANOVA, ***(P<0.001). Error bars represent SEM. Scale bar represents 10 µm. **(C)** Quantitative estimation of hyphal cells was performed in SC + 0.1% glucose, in the presence and absence of NPD10084 or PKM2-IN-6 at a concentration of 100 µM. Statistical analysis was conducted using one-way ANOVA, ***(P<0.001). Error bars represent SEM. Scale bar represents 10 µm. **(D)** Quantitative estimation of hyphal cells was performed in YP medium supplemented with 10% serum, in the presence and absence of NPD10084 or PKM2-IN-6 at a concentration of 100 µM. Statistical analysis was conducted using one-way ANOVA, ***(P<0.001). Error bars represent SEM. Scale bar represents 10 µm. **(E)** CFU assay was performed in morphogenesis-inducing media with and without 100 µM of NPD10084 or PKM2-IN-6. Statistical analysis was conducted using one-way ANOVA, ns (non-significant). Error bars represent SEM.

To determine whether the observed inhibition of hyphal differentiation by these compounds under various filamentation-inducing conditions was independent of effects on overall fungal growth, we performed CFU assays using the wild-type strain grown in the aforesaid media conditions, in the presence of 100 µM NPD10084 and PKM2-IN-6, while the untreated wild-type strain was used as a control. CFU analysis revealed no significant difference in *C. albicans* growth between inhibitor-treated and untreated wild-type controls across all of these media conditions (Fig 3E). These results indicate that, at the tested concentration, neither NPD10084 nor PKM2-IN-6 impairs fungal viability but specifically blocks fungal morphogenesis under all the tested filamentation-inducing conditions.

### Pyruvate Kinase Inhibition Induces Extensive Transcriptional Remodeling of Gene Networks Associated with Filamentation, Biofilm Formation, Antifungal Susceptibility and Virulence

Our previous results established that PKM2-IN-6 potently blocks hyphal differentiation across multiple morphogenesis-inducing media without affecting fungal viability. To elucidate the molecular mechanisms underlying PKM2-IN-6-mediated inhibition of hyphal differentiation, we performed comparative RNA-Sequencing to identify the transcriptional reprogramming associated with pyruvate kinase inhibition in *C. albicans* under morphogenesis-inducing condition. In order to do this, *C. albicans* wild-type was cultured under morphogenesis-inducing conditions (Spider medium with glucose) in the presence or absence of PKM2-IN-6 for 12 hours, followed by RNA isolation, sequencing, and differential expression analysis (Fig 4A). Volcano plot analysis of the RNA-Seq data revealed extensive, statistically significant transcriptional changes upon PKM2-IN-6 treatment, with a substantial number of genes showing both upregulation and downregulation (Fig 4B). This broad transcriptional shift indicates that pharmacological inhibition of pyruvate kinase substantially reshapes the transcriptional landscape of *C. albicans* during hyphal differentiation.

**Fig 4.**
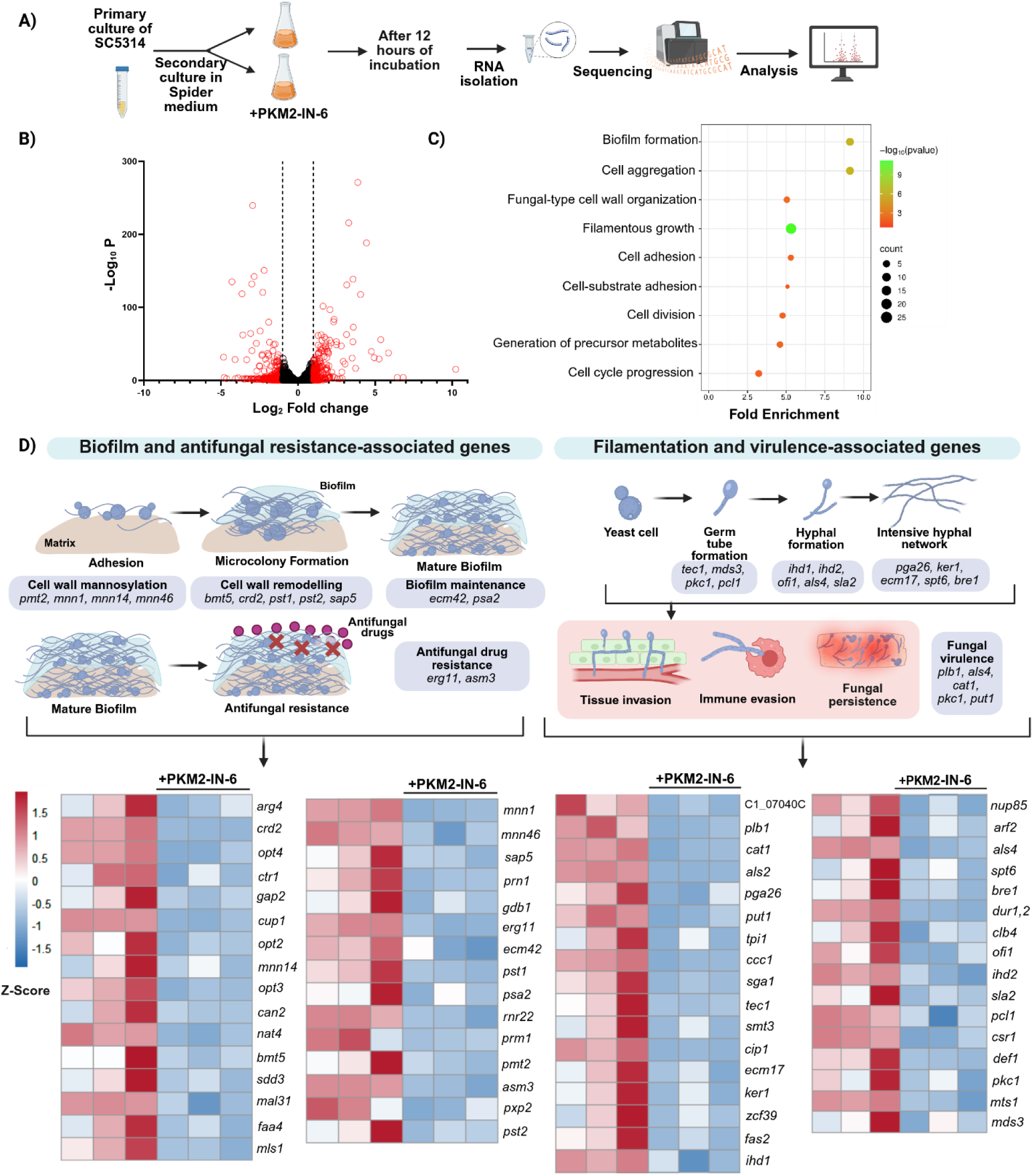
Pyruvate kinase inhibition induces extensive transcriptional remodeling associated with hyphal differentiation, biofilm formation and fungal virulence. (A) Schematic representation of the experimental workflow. *C. albicans* SC5314 cells were inoculated into Spider medium supplemented with glucose and cultured in the presence of the pyruvate kinase inhibitor (+PKM2-IN-6). Following 12 hours of incubation under filament-inducing conditions, total RNA was isolated and subjected to RNA-Sequencing and differential gene expression analysis. (B) Volcano plot showing differentially expressed genes following pyruvate kinase inhibition. Each dot represents a single gene, plotted according to its log₂ fold change (x-axis) and –log₁₀ adjusted P-value (y-axis). Genes with significant differential expression are distributed on both sides of the plot. (C) Gene Ontology (GO) enrichment analysis of downregulated genes. The x-axis indicates fold enrichment of each GO term. Bubble size represents the number of genes associated with each GO term, and bubble color represents enrichment significance (−log₁₀ adjusted P-value). (D) Heatmap illustrating expression profiles of genes associated with biofilm formation and antifungal resistance (left) and filamentation and virulence (right) following pyruvate kinase inhibition. Gene expression values are shown as Z-scores, where red indicates relatively higher expression and blue indicates relatively lower expression. The accompanying schematic summarizes the contribution of fungal morphogenesis to biofilm formation, antifungal resistance, tissue invasion, immune evasion, fungal persistence, and host morbidity.

To identify the biological processes most affected by PKM2-IN-6 treatment, differentially expressed genes were subjected to Gene Ontology (GO) enrichment analysis (Fig 4C). GO analysis demonstrated that the differentially expressed downregulated genes were predominantly associated with biological processes involved in fungal morphogenesis and virulence (Fig 4C). Among the most significantly enriched categories were genes involved in biofilm formation, filamentous growth, cell adhesion, fungal cell wall organization, cellular bud neck septin ring organization, and regulation of cell wall biogenesis. In addition, enrichment of genes related to the mitotic cell cycle, cell division, cellular metabolism, and generation of precursor metabolites suggested that pyruvate kinase inhibition broadly affects cellular physiology beyond just glycolytic metabolism.

Given the strong enrichment for biofilm-related genes in our GO analysis, we next examined the expression of genes known to function at discrete stages of biofilm development in *C. albicans* (Fig 4D, left). Genes involved in cell wall mannosylation and early adhesion (*pmt2, mnn1, mnn14, mnn46* (56, 57)), cell wall remodeling during microcolony formation (*bmt5, crd2, pst1, pst2, sap5* (58, 59)), and biofilm maintenance in the mature biofilm (*ecm42, psa2* (60, 61)) were all consistently downregulated in PKM2-IN-6-treated samples relative to untreated controls across biological replicates, as reflected in the corresponding Z-score heatmaps. Genes associated specifically with antifungal drug resistance in mature biofilms, including the ergosterol biosynthesis gene *erg11* and *asm3* (62–64), likewise showed significantly reduced expression upon PKM2-IN-6 treatment. Because *erg11* is a principal determinant of azole susceptibility and biofilm-associated drug tolerance, its downregulation suggests that PKM2 inhibition may compromise not only biofilm architecture but also the drug-resistant phenotype characteristic of mature biofilms (62, 63). We next assessed the expression of genes governing the sequential steps of filamentation, from germ tube formation through hyphal elongation to formation of an intensive hyphal network, as well as genes linked to tissue invasion, immune evasion, and fungal persistence during infection (Fig 4D, right). Germ tube-associated regulators (*tec1, mds3, pkc1, pcl1* (65–68)), hyphal formation genes (*ihd1, ihd2, ofi1, als4, sla2* (69–71)), and genes required for extensive hyphal network formation (*pga26, ker1, ecm17, spt6, bre1* (72–77)) were uniformly downregulated in PKM2-IN-6-treated cells. Similarly, genes linked to fungal virulence, tissue invasion, and immune evasion (*plb1, als4, cat1, pkc1, put1*(67, 70, 78–80)), as well as an additional list of filamentation/virulence-associated genes (*nup85, arf2, als4, spt6, bre1, dur1,2, clb4, ofi1, ihd2, sla2, pcl1, csr1, def1, pkc1, mts1, mds3*), showed the same pattern of reduced expression relative to untreated controls. The concomitant downregulation of genes spanning hyphal initiation (germ tube formation), progression (hyphal elongation) and hyphal network formation, together with biofilm formation and virulence, indicates that the inhibition of pyruvate kinase activity by PKM2-IN-6 broadly affects multiple transcriptional networks associated with fungal morphogenesis and virulence rather than affecting a single regulatory node.

Collectively, our data demonstrates that the inhibition of pyruvate kinase suppresses multiple gene networks required for fungal morphogenesis and pathogenicity. The coordinated downregulation of biofilm, antifungal resistance, and filamentation-associated genes supports a central role for pyruvate kinase in regulating virulence-associated transcriptional programs in *C. albicans*, providing a mechanistic basis for the impaired filamentation and pathogenic phenotypes observed upon pyruvate kinase inhibition (23).

### Pyruvate Kinase Inhibitors Disrupt Biofilm Formation and Enhance the Efficacy of Conventional Antifungal Drugs

The morphological transition from yeast to hyphal form is critical for biofilm formation in *C. albicans* (81). Our transcriptomic analysis indicated that pyruvate kinase inhibition broadly represses virulence-associated transcriptional programs in *C. albicans*, including pathways governing biofilm formation, cell adhesion, filamentous growth, and antifungal resistance. In particular, the downregulation of multiple biofilm-associated genes and the enrichment of GO terms related to biofilm development and fungal morphogenesis suggested that pyruvate kinase activity is required for the establishment of mature biofilms. Since hyphae constitute the structural framework of *C. albicans* biofilms, we next investigated whether these transcriptional alterations result in impaired biofilm formation following pharmacological inhibition of pyruvate kinase (Fig 5A). Our results demonstrate that treatment with both inhibitors showed a substantial reduction in biofilm formation, with a ∼77% decrease in biofilm formation, in the presence of NPD10084 and PKM2-IN-6 compared to the untreated controls (Fig 5B and 5C). This significant inhibition suggests that these compounds effectively impair the development of *C. albicans* biofilms, likely through their interference with the yeast-to-hyphae transition. Consequently, the inhibitors not only hinder morphological changes essential for virulence but also potentially enhance the susceptibility of *C. albicans* biofilms to antifungal therapies.

**Fig 5.**
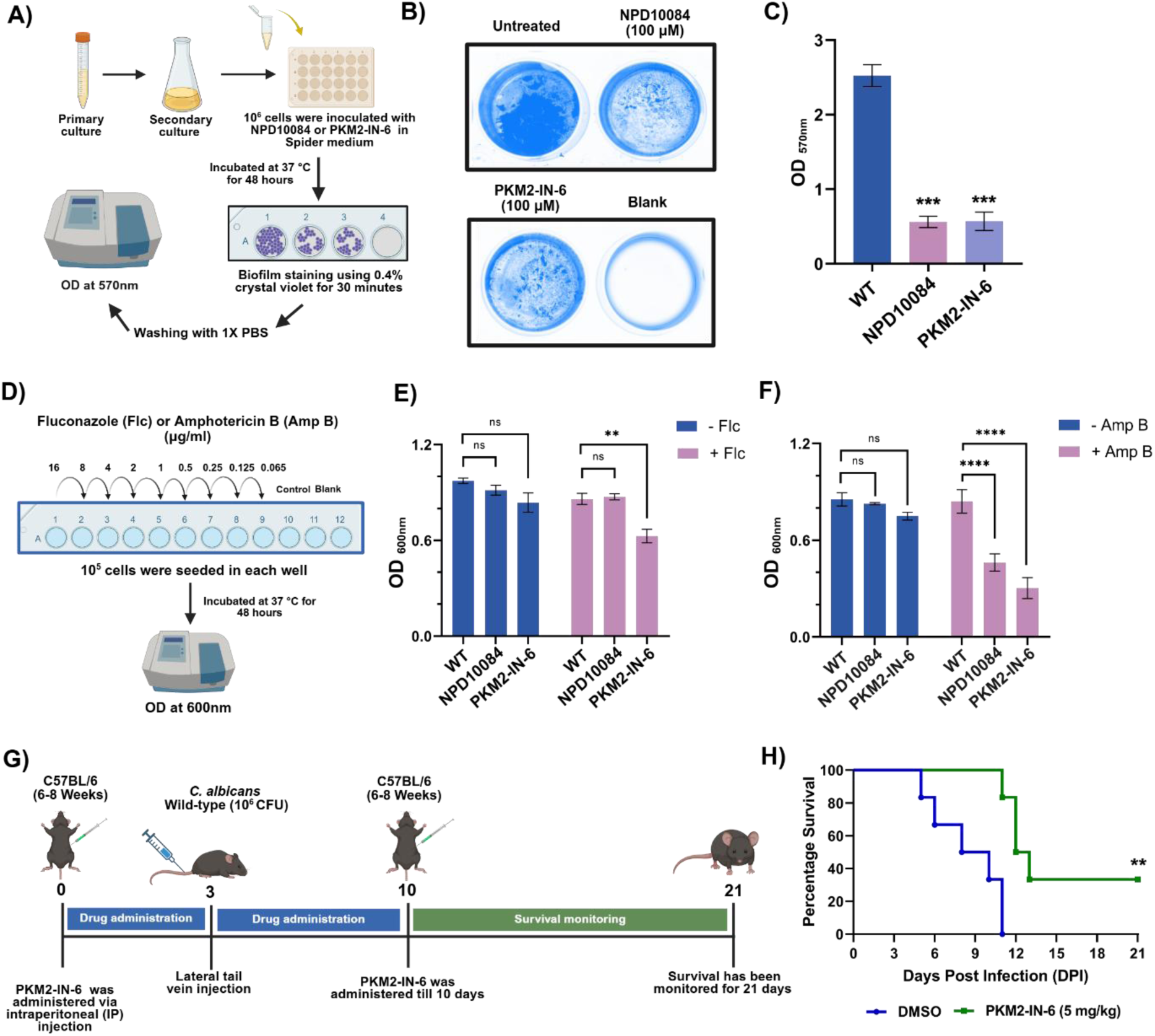
Pyruvate kinase inhibition disrupts biofilm formation, enhances antifungal efficacy and attenuates virulence in *C. albicans* by suppressing hyphal differentiation. **(A)** Schematic overview of the experimental procedure for biofilm assay. **(B)** A total of 10^6^ wild-type cells were cultured in Spider medium in the presence and absence of NPD10084 or PKM2-IN-6 at 100 µM. The cells were incubated at 37 °C for 48 hours and biofilm assay was performed by the crystal violet method. **(C)** Crystal violet-stained biofilm was quantified by measuring OD_570_. Statistical analysis was performed using one-way ANOVA, ***(P<0.001). Error bars represent SEM. **(D)** Schematic overview of the experimental procedure for the antifungal susceptibility test. **(E)** A total of 10^5^ wild-type cells were cultured in Spider medium in the presence and absence of fluconazole (Flc) at 0.25 µg/ml, combined with NPD10084 or PKM2-IN-6 at 100 µM. The cells were incubated at 37 °C for 48 hours, and growth was measured using a spectrophotometer at OD_600_. Statistical analysis was performed using one-way ANOVA, **(P<0.01) and ns (non-significant). Error bars represent SEM. **(F)** A total of 10^5^ wild-type cells were cultured in Spider medium in the presence and absence of amphotericin B (Amp B) at 0.065 µg/ml, combined with NPD10084 or PKM2-IN-6 at 100 µM. The cells were incubated at 37 °C for 48 hours, and growth was measured using a spectrophotometer at OD_600_. Statistical analysis was performed using one-way ANOVA, ****(P<0.0001) and ns (non-significant). Error bars represent SEM. **(G-H)** 6-8-week-old C57BL/6 mice were administered with PKM2-IN-6 (5mg/kg) for 10 days, following a schedule of two consecutive days of treatment with a one-day interval. On day 3, mice were intravenously challenged via the lateral tail vein with 10⁶ CFU of the *C. albicans*. Post-infection, animals were monitored daily for 21 days to assess survival, morbidity, and mortality. Survival percentages were statistically evaluated by the Log-rank Mantel-Cox test, **(P<0.01) (n=6).

Biofilm formation and fungal morphogenesis are closely linked processes that confer high-level resistance to multiple frontline antifungal drugs in *C. albicans*. Previous studies have demonstrated that downregulation of hyphae-associated and biofilm regulatory genes (e.g., *ras1, cyr1, efg1, ume6, hgc1, als3, hwp1, bcr1,* and *cdc28*) significantly increases the antifungal susceptibility in *C. albicans* (82, 83). In addition, deletion of *hog1* has been shown to compromise biofilm formation leading to enhanced sensitivity to antifungal agents (84, 85). Interestingly, *C. albicans* morphogenesis-defective mutants (*ΔΔefg1* and *ΔΔcph1*), which are impaired in biofilm formation and yeast-to-hyphae transition, exhibit increased antifungal susceptibility under both biofilm-inducing and non-inducing conditions, including enhanced sensitivity to amphotericin B compared to the wild-type strain (86). Specifically, the central regulator of morphogenesis, Efg1 plays a critical role in antifungal susceptibility in *C. albicans*, as its deletion increases sensitivity to ergosterol-targeting drugs by altering membrane composition and enhancing drug permeability (87). Moreover, several reports suggest that *C. albicans* morphogenesis can be blocked by conventional antifungal drugs such as fluconazole, ketoconazole, and amphotericin B (88–90). Taken together, these previous findings emphasize that morphogenesis and its regulatory networks are central determinants of antifungal resistance in *C. albicans*. Given that pyruvate kinase inhibitors effectively suppressed hyphal differentiation (Fig. 2A and 2B), caused broad transcriptional repression of genes involved in fungal morphogenesis, biofilm formation, antifungal resistance, and virulence (Fig. 4), and consequently impaired biofilm formation (Fig. 5A–5C), we sought to determine whether these inhibitors exhibit any synergistic effects when used in combination with well-known antifungal drugs and enhance their efficacy. To evaluate the combinatorial effects of pyruvate kinase inhibitors with conventional antifungals on inhibiting the growth of *C. albicans*, we tested different concentrations (16–0.065 µg/ml) of fluconazole and amphotericin B in Spider medium, with or without the pyruvate kinase inhibitors NPD10084 or PKM2-IN-6 (100 µM) (Fig 5D) and monitored the growth of *C. albicans* after 48 hours of treatment. At 0.25 µg/ml, fluconazole in combination with PKM2-IN-6 decreased growth by 27% relative to fluconazole alone, indicating a synergistic interaction, whereas NPD10084 showed no significant effect (Fig 5E). In contrast, in the presence of amphotericin B, PKM2-IN-6 and NPD10084 reduced growth by 64% and 45%, respectively, at 0.065 µg/ml compared to amphotericin B alone suggesting that the presence of the inhibitors enhanced fungal susceptibility towards amphotericin B compared to the control (Fig 5F). These findings suggest that pyruvate kinase inhibitors, particularly PKM2-IN-6, enhanced the efficacy of existing antifungals, including fluconazole and amphotericin B and highlight the potential of such combination therapies. Overall, the results indicate that inhibition of hyphal differentiation significantly increases antifungal susceptibility in *C. albicans*.

### PKM2-IN-6 Significantly Attenuate *C. albicans* Virulence in a Murine Model of Systemic Candidiasis

Fungal morphogenesis is a key virulence mechanism employed by *C. albicans* to establish successful infection in the host (16). Previous studies have demonstrated that glycolysis-defective mutant *ΔΔpyk1* (lacking pyruvate kinase) exhibits attenuated virulence in murine model of candidiasis. However, the mechanisms underlying this observed phenotype was not explored in that study (23). Since PKM2-IN-6 emerged as the most potent inhibitor of hyphal differentiation and transcriptomic analysis revealed coordinated repression of multiple genes associated with fungal morphogenesis and virulence in the presence of PKM2-IN-6, we next evaluated whether these effects translated into reduced pathogenicity in a murine model of systemic candidiasis. In order to do this, 6-8-week-old C57BL/6 mice were administered PKM2-IN-6 (5 mg/kg) via intraperitoneal injection for 10 days. On the third day post-administration, wild-type *C. albicans* (SC5314) (10^6^ cells) was injected through the lateral tail vein, and survival was monitored throughout the study period (Fig 5G). We observed that treatment of mice with PKM2-IN-6 resulted in significantly improved survival compared to the vehicle-treated mice (Fig 5H). These results suggest that targeting pyruvate kinase is a promising antifungal strategy for treating *C. albicans* infections.

### Comparative Docking and Interaction Analysis of identified Inhibitors with *C. albicans* Pyruvate Kinase

To gain the structural insights into the nature of interaction between the identified inhibitors and *C. albicans* pyruvate kinase (CaPyk1), *in silico* molecular docking analysis was carried out using the CaPyk1 pyruvate kinase protein structure (S2A Fig). Since no co-crystal structure of CaPyk1 was available in the PDB database and initial sequence or structure analysis did not confirm the most likely ligand-binding site of identified inhibitors, the blind docking was performed, followed by focused docking at the site-specific level for enrichment of binding poses. From blind Docking analysis, multiple binding sites were observed as depicted in figure 6, with population and docking energy at each site-specific cluster. The clustering of docked ligand conformations was based on binding modes and associated docking energies. From these clusters, the top five binding sites, i.e. site S1 to site S5, were selected based on their cluster ranking and distinct spatial locations (functionally relevant) on CaPyk1 (Fig 6A-6E). Lig. 1 (NPD10084, CID 29148165), and Lig. 2 (PKM2-IN-6, CID 148613955), revealed multiple binding sites (S1–S5) with distinct binding poses (Fig 6A and 6B) and for ATP in blind docking, multiple orientations were also observed at site S1. For Lig. 1, among the identified five sites across the protein, with Site-1 (S1; green) represents the most populated cluster, indicating strong site preference (Fig 6A). Other sites (S2–S5; orange, red, brown, and magenta) showed fewer conformers, suggesting comparatively lower affinity. Similarly, Lig. 2 also shows multiple sites (S1–S5), but displayed a more dispersed distribution, with conformers present at S4 and S5, indicating reduced site-specificity relative to Lig. 1 (Fig 6B), however, for both ligands, site S1 is preferable.

**Fig 6.**
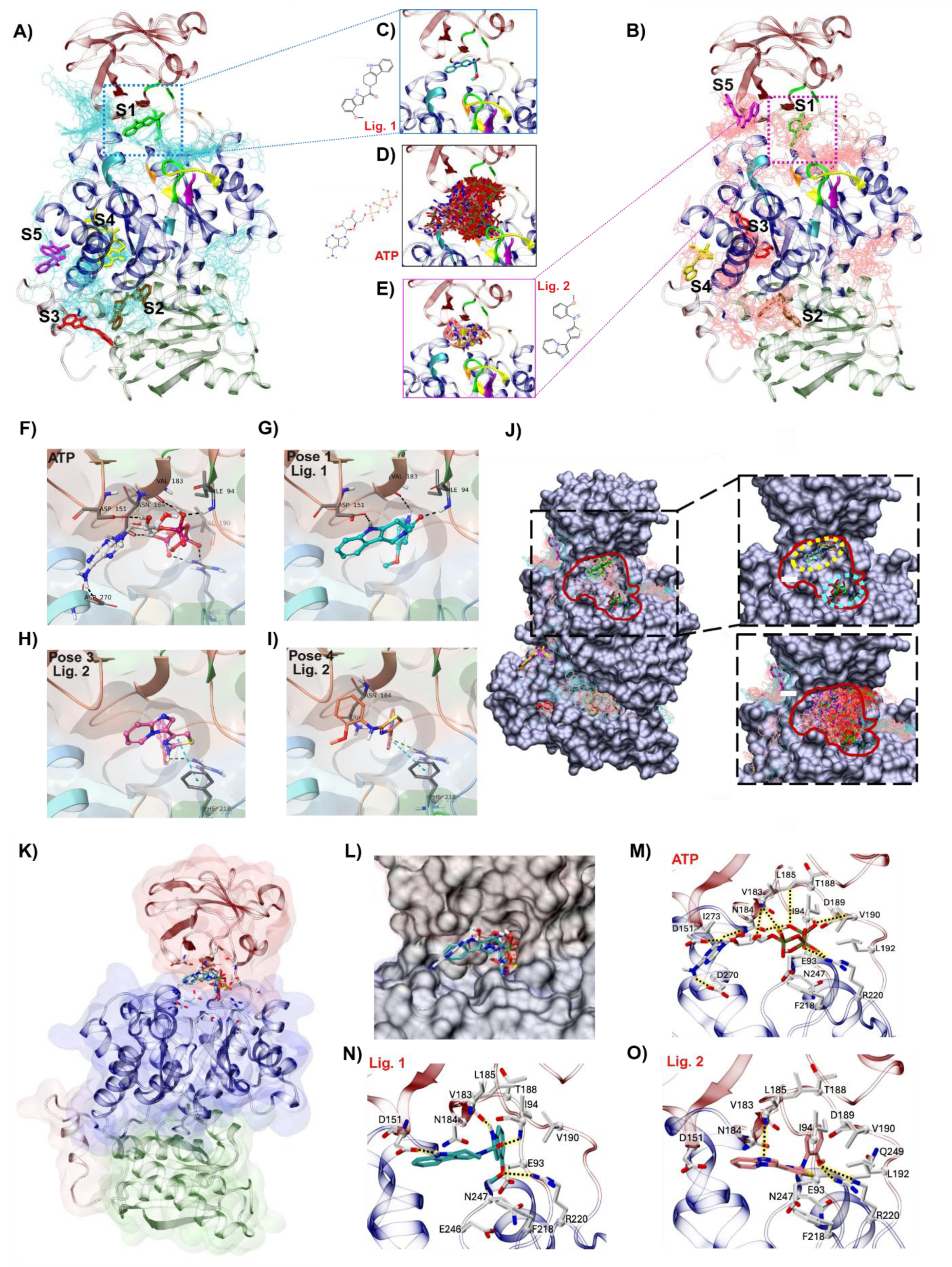
Comparative docking and interaction analysis of pyruvate kinase inhibitors with *C. albicans* pyruvate kinase (CaPyk1). **(A)** Identification and characterization of ligand-binding pockets in pyruvate kinase using blind docking of Lig. 1. **(B)** Identification and characterization of ligand-binding pockets in pyruvate kinase using blind docking of Lig. 2. **(C)** Detailed view of ligand interactions at Site-1, highlighting binding poses of Lig. 1. **(D)** Detailed view of ATP binding interactions at Site-1. **(E)** Detailed view of ligand interactions at Site-1, highlighting binding poses of Lig. 2. **(F)** Representative ATP binding pose illustrating critical interactions and conformational stability within the binding pocket. **(G)** Representative binding pose 1 of Lig. 1, showing key interactions and conformational stability within the binding pocket. **(H)** Representative binding pose 3 of Lig. 2 showing key interactions and conformational stability within the binding pocket. **(I)** Representative binding pose 4 of Lig. 2 showing key interactions and conformational stability within the binding pocket. **(J)** Surface-based analysis of ligand binding at Site-1 in pyruvate kinase, showing the overall surface with the Site-1 region highlighted (dashed box). Clustering of docked ligand poses is displayed alongside close-up views of the best docked ligands, including ATP within subpocket 1, the co-crystal ATP binding pose from superimposed *Leishmania mexicana* pyruvate kinase (PDB ID: 3HQP) within subpocket 2, and the distribution of blind docking poses indicating preferential ligand occupancy at subpocket 1. **(K)** Overall pyruvate kinase structure with the Site-1 binding pocket indicated (boxed). **(L)** Surface representation illustrating ligand occupancy within the Site-1 pocket. **(M)** Interaction map depicting key contacts of ATP at Site-1. **(N)** Binding interactions of Lig. 1 at Site-1. **(O)** Binding interactions of Lig. 2 at Site-1. Key interacting residues are depicted as sticks with dashed lines indicating hydrogen bonds.

Focused docking at S1 demonstrated that Lig. 1 adopts tightly clustered conformations within a single pocket, confirming stable and site-specific binding (Fig 6C). In contrast, Lig. 2 showed multiple conformers with slight orientation differences, suggesting weaker or less stable interactions (Fig 6E). ATP docking at S1 revealed multiple conformations within the same pocket, consistent with its functional flexibility as a natural substrate (Fig 6D). Binding energy analysis from blind docking showed that Lig. 1 and Lig. 2 exhibited energies ranging from −10.92 kcal/mol and −8.62 kcal/mol, respectively. Focused docking further confirmed strong binding at S1, with Lig. 1 showing an average docking energy of −9.14 kcal/mol (S2D Fig), Lig. 2 −7.7 kcal/mol (S2E Fig), and ATP ranging from −6.02 kcal/mol (S2C Fig).

To confirm the most likely poses of both ligands, the Binding pose meta dynamics (BPMD) analysis was carried out (Fig 6F-6I), where the graph shows ligand RMSD (Å) versus simulation time (ns) for Lig. 1 and Lig. 2 (S2B Fig). Pose-1 (in blue) exhibits the lowest RMSD throughout the simulation, indicating high stability. Pose-2 (green) and Pose-5 (magenta) show a significant rise in RMSD, suggesting poor stability and likely unfavorable binding. Pose-3 (red) and Pose-4 (cyan) display similar moderate RMSD values, indicating intermediate stability. From these observations, Pose-1 is the most stable, Pose-3 and Pose-4 are moderately stable, while Pose-2 and Pose-5 are unstable. We have also performed superimposition analysis of the ATP-bound pyruvate kinase from *Leishmania mexicana* (91) with our docked complexes and observed that the docked ATP and the best-selected poses occupied the same cavity in site-1 (Fig 6J). In contrast, superimposition showed that the ATP co-crystal occupies the same site S1 with a diverse binding pose. We identified that site-1 pocket comprises two distinct sub-pockets within the same site; hence, we termed them sub-pocket 1 and sub-pocket 2 based on the docked ATP and co-crystal ATP orientations. We have also shown the distribution of docked ATP multiple conformers within S1. This clearly indicates that the majority of conformers occupied both sub-pockets at S1 site, but an exact match to the co-crystal pose was not observed, possibly due to slight change in CaPyk1 with respect to crystal. Also, in our filtering criteria, the ATP docked to sub-pocket 2 is not among the top-ranked poses. This observation suggests that though the identified inhibitors and ATP occupy the same site, however, they bind to distinct sub-pockets within the same ATP-binding site, indicating a possible competitive mode of inhibition.

To underscore the mode of inhibition, a detailed interaction analysis is needed. Interaction fingerprinting showed that ATP established multiple hydrogen bonds and hydrophobic interactions with pocket residues (Fig 6K-6O). The main involving key residues with ATP are E93, I94, D151, V183, L185, T188, V190, L192, F218, R220, N247 and D270 (Fig 6M). Though binding slightly different poses, in the same S1 pocket, the Lig. 1 binding mode (Fig 6N) also attains fewer hydrogen bonds, which involve residues I94, D151, V183, and R220. Similar to Lig. 1, the same pocket was occupied by Lig. 2 as well (Fig 6O), however, it has shown considerably limited hydrogen bond interactions, mainly with V183 and R220. Overall, the binding pocket is stabilized by a combination of polar and hydrophobic interactions, with residues such as E93, L185, T188, N247, F218, and R220 playing key roles. Variations in ligand orientation and interaction profiles likely contribute to differences in binding affinity and specificity.

In addition to the docking energies of both ligands, the thermodynamic analysis was also calculated using the MM-GBSA approach. The results from this study yield the values that support the docking results. The ΔG calculated via MM-GBSA for Lig. 1 using the best docked pose is -42.47 kcal/mol with 148 conformers and for Lig. 2, it is -40.94 kcal/mol it is 99 conformers, respectively, confirming that both ligands occupy the S1 site, however, Lig. 1 seems to be a better inhibitor.

## Discussion

Fungal morphogenesis is a central determinant of *Candida albicans* pathogenicity, yet therapeutic strategies that directly target this process remain largely unexplored. The discovery of small-molecule inhibitors targeting morphogenesis has become a key approach for developing anti-virulence therapies against *C. albicans* (49, 92). Because filamentation is central to tissue invasion and biofilm formation, several groups have employed large-scale phenotypic screening of chemical libraries to discover compounds that selectively disrupt the yeast-to-hyphae transition without compromising the overall fungal growth (93). Such high-throughput efforts have successfully identified diverse bioactive molecules that suppress filamentation, demonstrating the feasibility and therapeutic promise of small-molecule based modulation of fungal virulence (92, 93). For example, a recent study identified a morphogenesis-specific inhibitor, filastatin, which significantly reduces *C. albicans* virulence in a mouse model of vulvovaginal candidiasis (VVC) (22). However, several studies including the aforementioned study have predominantly focused on targeting signaling and regulatory pathways, including the cAMP–PKA and HOG pathway, as well as transcription factors such as Efg1, Cph2, and Tec1, typically under serum-induced conditions (54, 93–96). Our previous work demonstrated that glycolysis is a critical regulator of fungal morphogenesis in *S. cerevisiae* and *C. albicans* and perturbation of glycolysis results in attenuated fungal virulence in *C. albicans* (25). Leveraging these findings, we performed a systematic phenotypic screening of a glycolysis compound lCbrary to identify small-molecule inhibitors glycolysis that effectively block hyphal differentiation in *C. albicans*. Our findings demonstrate that perturbation of glycolysis, particularly the late glycolytic step that results in the conversion of PEP to pyruvate, profoundly affects hyphal development without compromising the overall growth of *C. albicans*, thereby establishing glycolysis inhibition as a viable antifungal intervention strategy. Our screening of 112 compounds targeting distinct nodes of glycolysis and associated metabolic pathways revealed that multiple steps in central carbon metabolism influence filamentation. Notably, inhibitors of glucose transport, hexokinase activity, and other downstream glycolytic enzymes moderately reduced hyphal differentiation without compromising the overall growth. These observations indicate that nutrient acquisition and metabolic flux act not merely as bioenergetic drivers but also as regulatory signals that influence morphogenetic programs. This observation is consistent with prior studies demonstrating that metabolic state and carbon availability regulate key signaling pathways governing hyphal induction,, including cAMP–PKA pathway (20, 97, 98) and nutrient-sensing networks like the Target of Rapamycin (TOR) pathway, (99) and Cek1 mediated MAPK signaling pathway (54). Collectively, these pathways integrate nutrient availability, cellular energy status, and environmental cues to regulate filamentation.

A major finding of this study is that pharmacological inhibition of the late glycolytic enzyme pyruvate kinase, particularly by NPD10084 and PKM2-IN-6, resulted in a significant reduction in hyphal differentiation under various filamentation-inducing conditions. Importantly, both of these compounds have been characterized in cancer systems, offering valuable mechanistic insights into the functional consequences of human pyruvate kinase PKM2 inhibition beyond its canonical metabolic role. NPD10084 exhibits potent antitumor activity in HCT116 xenograft models by selectively targeting the non-glycolytic functions of PKM2. It disrupts PKM2 interactions with key transcriptional regulators, including β-catenin and STAT3, thereby attenuating downstream oncogenic signaling pathways associated with cell proliferation, survival, and tumour progression, underscoring the role of PKM2 as a critical signaling hub in cancer (29). Similarly, PKM2-IN-6 (compound 7d) is a rationally designed inhibitor that targets cancer metabolism via PKM2. In the 4T1 allograft model of triple-negative breast cancer in immunocompetent mice, it significantly reduced tumor burden, demonstrating robust in vivo efficacy. Notably, its low toxicity toward normal cells suggests a favorable therapeutic index and selective targeting of tumor-associated metabolic and signaling pathway (30). Notably, in our system, the inhibitory effect of NPD10084 and PKM2-IN-6 on hyphal differentiation occurred without any significant reduction in fungal viability, indicating selective interference with fungal morphogenesis rather than general cytotoxicity. The inability of exogenous sodium pyruvate to rescue the filamentation defect suggests that hyphal inhibitory effects exerted by NPD10084 and PKM2-IN-6 are not due to simple depletion of the downstream metabolite pyruvate, but likely due to the disruption of overall glycolytic flux and glycolysis-dependent metabolic and signaling processes. Impaired pyruvate kinase activity likely reduces glycolytic flux, leading to the accumulation of PEP and upstream intermediates, thereby perturbing carbon flux and cellular homeostasis (46). Furthermore, elevated PEP levels may inhibit upstream enzymes such as TPI, further restricting glycolytic activity (48). Together, these findings identify pyruvate kinase as a critical metabolic node of glycolysis that is important for fungal morphogenesis in *C. albicans* and supports targeted intervention of metabolic pathways that are critical for virulence-associated processes like hyphal differentiation as a promising antifungal therapeutic strategy.

Morphogenetic switching is a fundamental determinant of *C. albicans* pathogenicity, and has been shown to be an important pre-requisite for multiple virulence-associated processes including adhesion, filamentation, biofilm formation, tissue invasion, immune evasion, and antifungal resistance (16,78). Our transcriptomic analysis demonstrated that inhibition of pyruvate kinase results in widespread transcriptional reprogramming that extends far beyond central carbon metabolism, indicating that pyruvate kinase functions as an important regulator of *C. albicans* pathogenesis. Rather than selectively affecting the expression of individual virulence factors, pyruvate kinase inhibition altered the expression of genes involved in cell adhesion, fungal-type cell wall organization, filamentous growth, biofilm formation, cell division, and metabolic precursor generation. These findings suggest that perturbation of glycolytic flux disrupts an integrated regulatory network that coordinates morphogenesis-associated fungal virulence. Transcriptomic profiling revealed that pyruvate kinase broadly regulates the transcriptional network required for hyphal morphogenesis. Hyphal differentiation in *C. albicans* requires coordinated activation of signaling pathways controlling germ tube emergence, polarized growth, cell wall remodeling, and maintenance of hyphal integrity (50, 100). Inhibition of pyruvate kinase suppressed the expression of genes associated with early morphogenetic signaling, including *tec1, mds3, pkc1* and *pcl1*, while simultaneously reducing the expression of hypha-associated genes such as *ihd1, ihd2, ofi1* and *als4*, which contribute to hyphal differentiation, adhesion, and polarized growth (65–71). Genes involved in maintaining hyphal architecture, including *pga26, ker1, ecm17, spt6* and *bre1* were also repressed, suggesting that pyruvate kinase influences not only hyphal initiation but also the maintenance of filamentous growth (72–75, 77). One of the most striking observations was the coordinated repression of genes involved in multiple stages of biofilm development. Successful biofilm formation requires sequential events including initial adhesion, cell wall remodeling, microcolony formation, extracellular matrix production, and biofilm maturation (101, 102). Pyruvate kinase inhibition reduced the expression of genes involved in cell wall glycosylation and mannan biosynthesis, including *bmt5, pmt2, mnn1, mnn14* and *mnn46*, which are required for maintaining cell wall architecture and efficient adhesion to biotic and abiotic surfaces (56, 57). In addition, genes associated with cell wall remodeling and biofilm structural integrity, including *crd2, pst1* and *pst2*, together with the hypha-associated secreted protease Sap5, were also downregulated, suggesting impaired microcolony development and biofilm organization (58, 59). Furthermore, repression of *ecm42* and *psa2*, along with decreased expression of antifungal resistance-associated genes such as *erg11* and *asm3*, indicates that pyruvate kinase activity is also important for maintaining mature biofilm physiology and the intrinsic drug tolerance associated with biofilm growth (60–64). Consistent with these molecular signatures, treatment with both pyruvate kinase inhibitors (NPD10084 and PKM2-IN-6) significantly reduced biofilm formation in vitro. Additionally, the hyphal morphotype of *C. albicans* exhibits greater resistance to antifungal agents compared to the yeast form (104, 105). This enhanced resistance to antifungal agents is attributed to biofilm production, change in the cell wall composition or membrane fluidity during the yeast-to-hyphae transition (106, 107), and due to the shared regulatory signaling pathways (108, 109). In *C. albicans*, the cAMP-PKA signaling pathway plays a key role in antifungal resistance. Deletion of *cdc35* (encoding adenyl cyclase), which results in reduced intracellular cAMP levels, impairs hyphal differentiation and consequently increases susceptibility to fluconazole. (110, 111). In addition to the cAMP-PKA pathway, upregulated expression of efflux pumps including MDR1 and CDR1/CDR2, in the hyphal cells also provides antifungal resistance in *C. albicans* (81, 112). Our data demonstrated that the presence of pyruvate kinase inhibitors significantly enhanced the efficacy of antifungal drugs, including fluconazole and amphotericin B. Notably, PKM2-IN-6 exhibited a synergistic effect with fluconazole, while both PKM2-IN-6 and NPD10084 markedly improved the efficacy of amphotericin B, highlighting the potential of combining fungal morphogenesis inhibitors with conventional antifungal drugs. As our experiments were performed in Spider medium under static conditions, which is ideal for biofilm formation, the observed synergistic effect in the presence of PKM2-IN-6 and NPD10084, could be due to the inhibitory effects exerted by the compounds on hyphal differentiation and biofilm formation under these conditions. In a recent study, a morphogenesis inhibitor, beauvericin, which targets efflux pumps and the TOR pathway, was shown to exhibit synergism in attenuating *C. albicans* infection when used in combination with azoles (107, 113). These findings support the concept that inhibition of morphogenesis can increase sensitivity to antifungals by impairing morphogenesis-associated resistance mechanisms. Importantly, combination therapy offers a dual advantage by targeting both fungal growth and virulence traits, enhancing therapeutic efficacy. Moreover, such strategies may also reduce the likelihood of resistance development, as they impose lower selective pressure compared to fungicidal monotherapies. Given the increasing resistance to frontline antifungals, including azoles and echinocandins (114), our results underscore the potential of metabolism-targeted combination approaches for improved antifungal intervention.

Since sustained hyphal growth is indispensable for host tissue penetration and biofilm maturation, these findings provide a molecular basis for the severe filamentation defects observed following pyruvate kinase inhibition. Consistent with the close relationship between morphogenesis and pathogenicity, pyruvate kinase inhibition also suppressed several established virulence determinants involved in host adaptation. Reduced expression of *plb1*, a major phospholipase required for tissue invasion (78), together with repression of *als4*, which mediate host adhesion (70), *cat1*, which protects against oxidative stress (79), and *put1* and *pkc1*, which contribute to metabolic adaptation and cell wall integrity (67, 80), collectively suggests that inhibition of pyruvate kinase compromises multiple virulence mechanisms required for successful host colonization and persistence. Rather than regulating individual pathogenicity genes independently, pyruvate kinase appears to coordinate a broad transcriptional network that integrates morphogenesis, stress adaptation, and virulence. Corroborating these observations, pharmacological inhibition of pyruvate kinase significantly improved host survival in a murine model of systemic candidiasis, demonstrating that inhibition of pyruvate kinase-dependent metabolic programs effectively attenuates virulence in vivo.Our observations are consistent with an earlier report demonstrating that the glycolysis-defective *ΔΔpyk1* strain exhibits attenuated virulence in mice (23). Although pyruvate kinase has traditionally been regarded as the terminal rate-limiting enzyme of glycolysis, increasing evidence suggests that metabolic enzymes function as critical regulators of cellular signaling and transcriptional responses. In *C. albicans*, central carbon metabolism is closely integrated with nutrient-responsive pathways such as the cAMP-PKA, TOR, and Cek1 MAPK pathways, which collectively govern hyphal development, biofilm formation, and virulence gene expression (20,54,75–77). Inhibition of pyruvate kinase is expected to reduce glycolytic flux while altering intracellular concentrations of phosphoenolpyruvate, pyruvate, ATP, and other metabolic intermediates, thereby disrupting cellular energy homeostasis and nutrient signaling. These metabolic perturbations are likely to indirectly influence signaling networks that regulate morphogenesis and pathogenicity. Although the precise molecular mechanism linking pyruvate kinase to these signaling pathways remains to be elucidated, our findings strongly support a model in which pyruvate kinase acts as a metabolic hub coupling glycolytic activity to the transcriptional programs governing biofilm formation, filamentation, antifungal resistance, and virulence. Overall, our study expands the functional role of pyruvate kinase beyond its established metabolic function and identifies it as a central regulator of the pathogenic lifestyle of *C. albicans*. The coordinated repression of genes governing adhesion, cell wall remodeling, hyphal differentiation, biofilm formation, antifungal resistance, and virulence provides a mechanistic framework for the broad phenotypic defects observed upon pyruvate kinase inhibition and highlights this enzyme as a promising target for the development of antifungal strategies that simultaneously attenuate multiple virulence attributes.

To gain structural insights into the mechanisms underlying inhibition of pyruvate kinase by NPD10084 and PKM2-IN-6, we performed *in silico* molecular docking analyses to identify potential ligand-binding sites within *C. albicans* CaPyk1. Comparative structural alignment with the ATP-bound pyruvate kinase from *Leishmania mexicana* revealed that the predicted ATP binding poses in CaPyk1 not exactly overlap with the canonical co-crystal ATP binding site observed in the reference structure. Instead, docking results indicated that ATP preferentially occupies an adjacent sub-pocket within the broader site-1 region, suggesting structural divergence in nucleotide binding architecture between these enzymes. Blind and focused docking identified multiple potential ligand-binding regions i.e. site S1-to-site S5, however, site S1 showing the most likely binding site in terms of docking energy and high population of conformers. At S1 site the BPMD followed by fee energy via MM-GBSA, confirm the most appropriate pose of both ligands. The poses of both ligands occupied the same sub-pocket preferred for ATP docking in CaPyk1, but distinct from the canonical catalytic ATP site in the reference enzyme. This suggests that the binding likely occurs at an adjacent regulatory region rather than the active site, consistent with a competitive mode of inhibition. Collectively, these findings provide strong evidence that our compounds likely exert their morphogenesis inhibiting effects by modulating the activity of the *C. albicans* pyruvate kinase enzyme.

In conclusion, our systematic screening of a glycolysis-targeting small-molecule library identified two promising inhibitors of pyruvate kinase that robustly block hyphal differentiation across diverse filamentation-inducing conditions without impairing fungal viability. Comparative RNA-Seq analysis further revealed that pyruvate kinase inhibition induces extensive transcriptional reprogramming characterized by coordinated repression of gene networks governing hyphal development, biofilm formation, cell adhesion, antifungal resistance, and fungal virulence, providing a mechanistic basis for the observed phenotypes. These inhibitors enhance the antifungal activity of conventional drugs through synergistic interactions by disrupting morphogenetic switching and biofilm formation, thereby increasing the sensitivity of *C. albicans* to these antifungal agents. Notably, treatment with PKM2-IN-6 also improved survival of the host in a murine model of systemic candidiasis, underscoring its therapeutic potential. Over the past several decades, only a limited number of antifungal drug classes have been developed, with azoles remaining the widely used class of antifungal agents (115–117). The increasing prevalence of resistance to these aforesaid antifungal drugs among various fungal pathogens highlights the urgent need for novel therapeutic strategies. In this context, glycolysis represents a promising target as it has been shown to be critical for fungal morphogenesis in multiple fungal species (25, 118–122), and our current study demonstrates that its perturbation selectively impairs morphogenesis, without affecting overall fungal growth. Taken together, these findings establish glycolysis as a promising, previously underexplored target for antifungal therapeutic intervention and supports targeting of key metabolic regulatory nodes of morphogenesis as a rational anti-virulence strategy.

## Materials and Methods

### Yeast Strains

A prototrophic diploid strain of *Candida albicans* SC5314 was used in this study.

### Media and Growth Conditions

YPD broth containing 10 g/l yeast extract, 20 g/l peptone, and 2% (w/v) glucose was used for the overnight growth of the cultures. For filamentation assays, Spider medium (nutrient broth 10 g/l, K_2_HPO_4_ 2 g/l, and 2% (w/v) glucose), RPMI medium (RPMI 1640, 165 mM MOPS, pH-7), Lee’s medium pH 6.8 (NH_4_)_2_SO_4_ 5 g/l, MgSO_4_.7H_2_O 0.2 g/l, K_2_HPO_4_ (Anhydrous) 2.5 g/l, NaCl 5 g/l, Glucose 12.5 g/l, alanine 0.5 g/l, leucine 1.3 g/l, lysin 1 g/l, methionine 0.1 g/l, ornithine 70 mg/l phenylalanine 0.5 g/l, proline 0.5 g/l, threonine 0.5 g/l, biotin 1 mg/l), SC medium (Yeast nitrogen base) + 0.1% glucose and YPD + 10% serum were used. For Biofilm, Difco Sabouraud Dextrose Broth and Spider medium were used. All filamentation assays were performed in 96-well plates, which were incubated at 37 °C for 48 hours. Biofilm assay was performed in a 24-well plate, which was incubated at 37 °C for 48 hours under static conditions.

### Filamentation Assay

For screening the glycolysis compound library, 10^5^ cells of *C. albicans* SC5314 (wild-type) were seeded in a 96-well plate containing Spider medium and one unique glycolysis inhibitor compound at 250 µM concentration from the ‘Glycolysis Compound Library’. For the compounds that exhibited growth defects at 250 µM, filamentation assays were performed at 100 µM and 10 µM. To assess filamentation under differential media conditions, we used Lee’s medium, RPMI medium, SC + 0.1% glucose, and YP + 10% serum, and inoculated with 10^5^ cells and 100 µM compounds. The 96-well plate was then incubated at 37 °C for 48 hours. After the incubation, cells were fixed with 4% paraformaldehyde (PFA), and single cells were imaged using a Zeiss Apotome microscope. The length-to-width ratio of individual cells was measured using ImageJ, and the percentage of hyphal cells in the total population was determined. Cells with a length-to-width ratio of 4.5 or greater were considered hyphal cells, as described previously (123). Statistical analysis was assessed by one-way ANOVA using GraphPad Prism (version 9.0). Error bars represent SEM. All the experiments were performed independently three times.

### Colony Forming Units (CFU) Assay

To determine the effect of the two selected inhibitors (NPD10084 and PKM2-IN-6) on general growth, we performed a CFU plate assay. 10^5^ cells of *C. albicans* SC5314 were seeded in a 96-well plate containing the two inhibitors at a 100 µM concentration in different morphogenesis inducing media. The 96-well plate was then incubated at 37 °C for 48 hours. After the incubation, cells were serially diluted, and 100 µl of cells were plated on YPD agar plates. These plates were incubated at 30 °C for 24 hours, and colonies were counted to calculate the number of colonies per milliliter. Statistical analysis was assessed by one-way ANOVA using GraphPad Prism (version 9.0). Error bars represent SEM. All statistical comparisons were based on a minimum of three independent biological replicates.

### Pyruvate Add-back Assay

*C. albicans* SC5314 cells (10⁵ cells per well) were seeded into 96-well plates containing Spider medium supplemented with 2 mM NaPy and treated with either NPD10084 or PKM2-IN-6 at a final concentration of 100 µM. Plates were incubated at 37 °C for 48 hours to induce filamentation. After incubation, cells were fixed with 4% PFA, and filamentation was quantified by single cell imaging in the presence and absence of sodium pyruvate.

### RNA-Sequencing

#### Sample Collection

Overnight primary culture of *C. albicans* SC5314 was inoculated into YPD broth, followed by a secondary culture in Spider medium containing PKM2-IN-6 at a concentration of 100 µM and cultures were incubated at 37 °C for 12 hours with shaking conditions.

#### RNA-Extraction

Total RNA was isolated from cells harvested after 12 hours of incubation using the hot acid-phenol method. Following a DNAse treatment to eliminate any genomic DNA from the sample, the extracted RNA was submitted to the CCMB next-generation sequencing facility for RNA-Seqencing. The NovaSeq 6000 equipment was used to sequence transcriptomes.

#### RNA-Sequencing Analysis

Raw sequencing reads were obtained in FASTQ format. The reference genome and annotation files for the *C. albicans* strain SC5314 were downloaded from the Candida Genome Database (CGD). Quality assessment of the raw reads was performed using FastQC (version 0.12.1). Adapter sequences were trimmed using Cutadapt (version 4.6). The reference genome was indexed with Hisat2 (version 2.2.1), and Samtools (version 1.20) was used to filter out multi-mapped reads and convert SAM files to BAM format. Read counting for uniquely mapped reads was done with FeatureCounts, using gene annotation data from the reference *C. albicans* SC5314 genome. Normalization and differential gene expression analysis were performed using R (version 4.3.2). Genes with a log2 fold change of ≥ 1 were considered to be upregulated, indicating at least a doubling in expression under the given condition, while those with a log2 fold change of ≤ -1 were considered to be downregulated.

### In vitro Biofilm Assay

To evaluate the anti-biofilm efficacy of pyruvate kinase inhibitors (NPD10084 and PKM2-IN-6) against *C. albicans*, we performed the crystal violet staining assay as previously described (124). Briefly, an overnight primary culture of *C. albicans* SC5314 was inoculated into YPD broth, followed by a secondary culture in Sabouraud Dextrose Broth. A total of 10^6^ cells were resuspended in 1 ml of Spider medium and seeded into 24-well microtiter plates. Pyruvate kinase inhibitors NPD10084 and PKM2-IN-6 were added to their respective wells at a concentration of 100 µM, and plates were incubated at 37 °C for 48 hours without shaking to allow biofilm formation. Following incubation, non-adherent planktonic cells were removed by washing the wells gently with sterile 1× PBS. Sessile biofilm cells attached to the well bottoms were stained with 0.4% crystal violet for 30 minutes. Excess stain was removed by washing with sterile 1× PBS. After air-drying the plates for 15 minutes, the bound crystal violet was solubilized in 96% ethanol. The absorbance of the dissolved stain, proportional to biofilm biomass, was measured spectrophotometrically at 570 nm.

### Antifungal Susceptibility Testing

To check the efficacy of antifungal drugs, including fluconazole and amphotericin B, against *C. albicans* in the presence and absence of NPD10084 and PKM2-IN-6, we performed standard MIC assays. For fluconazole and amphotericin B, we used a range of 16 µg/ml to 0.065 µg/ml. For this assay, we seeded 10^5^ cells per well with or without NPD10084 and PKM2-IN-6, along with fluconazole and amphotericin B, with appropriate controls. The plate was then incubated at 37 °C for 48 hours. After incubation, absorbance was measured at 600 nm, and a graph was plotted using sub-inhibitory concentrations of fluconazole and amphotericin B. Statistical analysis was assessed by one-way ANOVA using GraphPad Prism (version 9.0). Error bars represent SEM. All experiments were performed in three independent biological replicates.

### In vivo Murine Model of Systemic Candidiasis

To study the in vivo antifungal activity of these pyruvate kinase inhibitors, we tested these compounds in a murine model of systemic candidiasis. A dose of 5 mg/kg of PKM2-IN-6 was administered to 6–8-week-old C57BL/6 mice via intraperitoneal (IP) injection for 10 days, following a regimen of two consecutive treatment days of compounds followed by a one-day interval. On day 3, mice were intravenously infected via the lateral tail vein with *C. albicans* wild-type strain SC5314 (10⁶ CFU). Following infection, mice were monitored daily for survival, as well as clinical signs of morbidity and mortality, for a period of 21 days. Survival data were analyzed with GraphPad Prism (version 9.0), and significance was calculated using the Log-rank Mantel-Cox test.

### Data curation for Molecular docking

We have retrieved the AlphaFold model of *C. albicans* pyruvate kinase (UniProt ID: P46614, 504 amino acids) and ligands (Lig. 1-NPD10084, PubChem ID: 29148165; Lig. 2-PKM2-IN-6, PubChem ID: 148613955) for docking.

### Docking of Small Molecules

Docking studies were carried out using AutoDock4 with AutoDockTools 1.5.7 (125). The modeled protein structure was prepared using the default protocol for the addition of missing atoms and hydrogens, followed by the assignment of Kollman charges to the protein. Ligands were prepared using the default parameters. Initially, blind docking was carried out by defining the whole protein as the grid center, with the number of grid points in the x, y, and z directions set to 130, 130, and 150, respectively. Docking was performed using genetic algorithms as the search parameters, with default docking run options and the Lamarckian GA for output docking results. GA runs were set for 200 in our study.

For focused docking, similar docking parameters were used, but grid centers were defined for each selected ligand at their respective sites (S1–S5) for both the ligand and ATP. Post-docking analysis was carried out using PyMOL and Schrödinger’s Maestro for visualization and post-processing. VMD was used to prepare figures in this study.

### Binding Pose Metadynamics (BPMD) Analysis

BPMD analysis was carried out using Schrödinger’s Desmond to identify the most stable binding poses of Lig. 1 and Lig. 2 among the predicted poses at site 1. BPMD assesses pose stability by performing metadynamics-based molecular dynamics simulations, in which the ligand RMSD (Root Mean Square Deviation) relative to its initial pose is used as the collective variable after binding-site superimposition to account for system drift. This RMSD is monitored throughout the simulation. Pose stability is evaluated based on the magnitude of ligand RMSD fluctuations, as well as the persistence of key interactions with the receptor and the surrounding environment. To improve statistical reliability, multiple independent simulations (10 × 10 ns for each pose) were performed, and the results were averaged. The resulting output showcases the average value of the collective variable over time, along with the Pose Score (lower values indicate greater stability) and PersScore (higher values indicate greater stability) (126).

### Thermodynamics Study

Following the docking analysis, the identified poses were subjected to MM-GBSA (Molecular Mechanics Generalized Born Surface area) calculations using Schrödinger’s Prime-MM-GBSA. This approach estimates the approximate binding free energy of each pose against the target receptor, enabling us to validate the most energetically favorable conformation within the protein pocket. For this analysis, we used both blind-docking and focused-docking poses as input ligands, with the prepared protein serving as the receptor. Each energy contribution is calculated by using the energies from the force field (OPLS4, (127) and the implicit solvation model.

The binding free energy in MM-GBSA is typically calculated as:

ΔG_“bind” (MM-GBSA) = G_“complex” - (G_“protein” + G_“ligand”)

Where: G_“complex” = Free energy of the protein-ligand complex;

G_“protein” = Free energy of the isolated protein;

G_“ligand” = Free energy of the isolated ligand

## Illustrations

Figure illustrations were created using Biorender (https://app.biorender.com).

## Animal Ethics Statement

All animal experiments in this manuscript were reviewed and approved by the Institutional Animal Ethics Committee (IAEC) of CSIR-Centre for Cellular and Molecular Biology (Approval number: 74/2024).

## Data Availability

All relevant data are within the paper and its supporting information files.

## Acknowledgements

We sincerely acknowledge the support and access to facilities provided by the CSIR–Centre for Cellular and Molecular Biology (CCMB), including the Fine Biochemicals Facility and Animal House Facility. SV gratefully acknowledges financial support from the Indian Council of Medical Research (ICMR) (IIRPSG-2024-01-02717), the Anusandhan National Research Foundation (ANRF) (SRG/2023/000470), and the Council of Scientific and Industrial Research (CSIR) (FTT070505). SA acknowledge the Department of Biotechnology (DBT) for funding through the National Network Project (NNP-BT/PR40189/BTIS/137/50/2022).

## Supporting Information

**S1 Fig.**
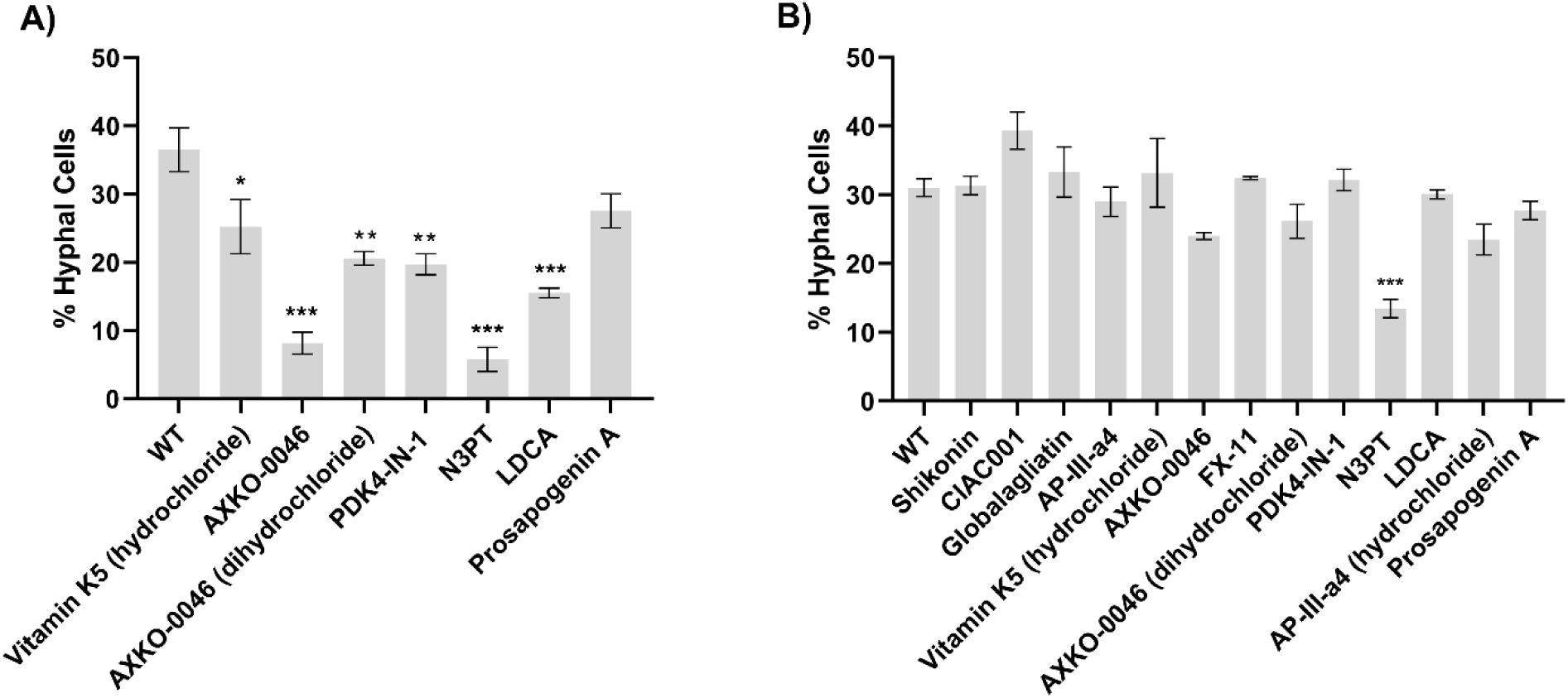
Effect of lower concentrations of glycolysis compounds that exhibit growth defects at higher concentrations on hyphal differentiation in *C. albicans*. Quantitative estimation of hyphal cells in the presence and absence of glycolysis inhibitors at 100 µM **(A)** and 10 µM **(B)** concentration with wild-type. A total of 10^5^ cells of the wild-type strain were cultured in Spider medium with and without the glycolysis inhibitors at 37 °C for 48 hours. Cells were fixed and imaged using Zeiss Apotome microscope. The length-to-width ratio of individual cells was measured using ImageJ, and the percentage of hyphal cells in the total population was determined. Cells with a length-to-width ratio of 4.5 or greater were considered hyphal cells. More than 300 cells were counted for each inhibitor. Statistical analysis was done using one-way ANOVA, ***(P<0.001), **(P<0.01) and *(p<0.1). Error bar represents SEM.

**S2 Fig.**
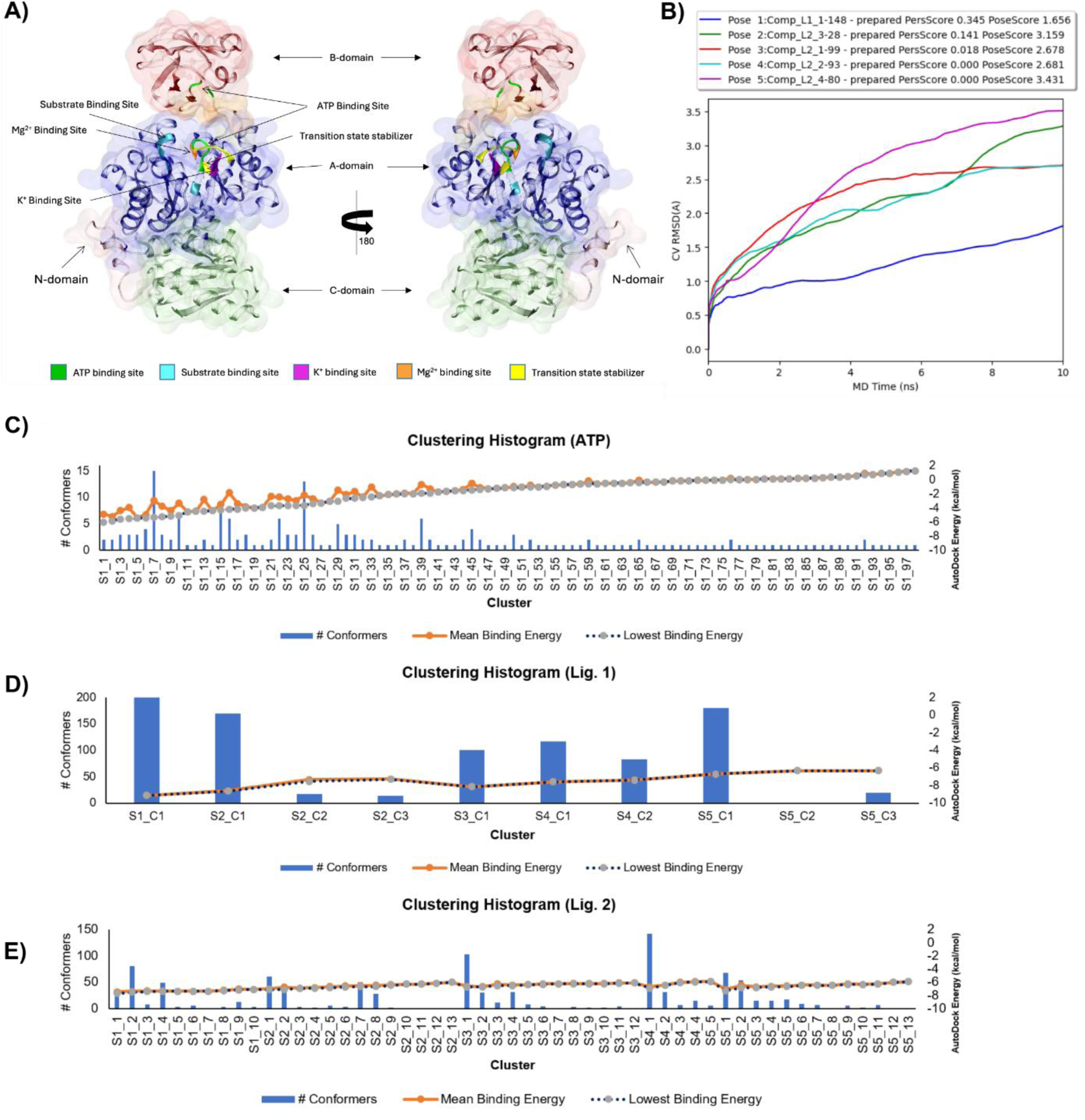
The structural organization and 3D orientation of the CaPyk1 and binding energy analysis for blind and focused docking of Lig. 1 and Lig. 2. **(A)** Structural organization and 3D orientations of pyruvate kinase illustrating the ATP and substrate binding sites. The protein is composed of N, A, B, and C domains, shown in distinct colors. Two different views are presented to highlight the spatial arrangement of domains and the positioning of the binding sites. **(B)** Time evolution of collective variable RMSD (CV-RMSD) for the selected docked poses **(C)** Clustering histogram of ATP docking poses showing the number of conformers per cluster, mean binding energy, and lowest binding energy at site-1. **(D)** Clustering histogram of Lig. 1 docking poses across predicted binding sites (S1–S5). **(E)** Clustering histogram of Lig. 2 docking poses across predicted binding sites (S1–S5).

**S1 Table:**
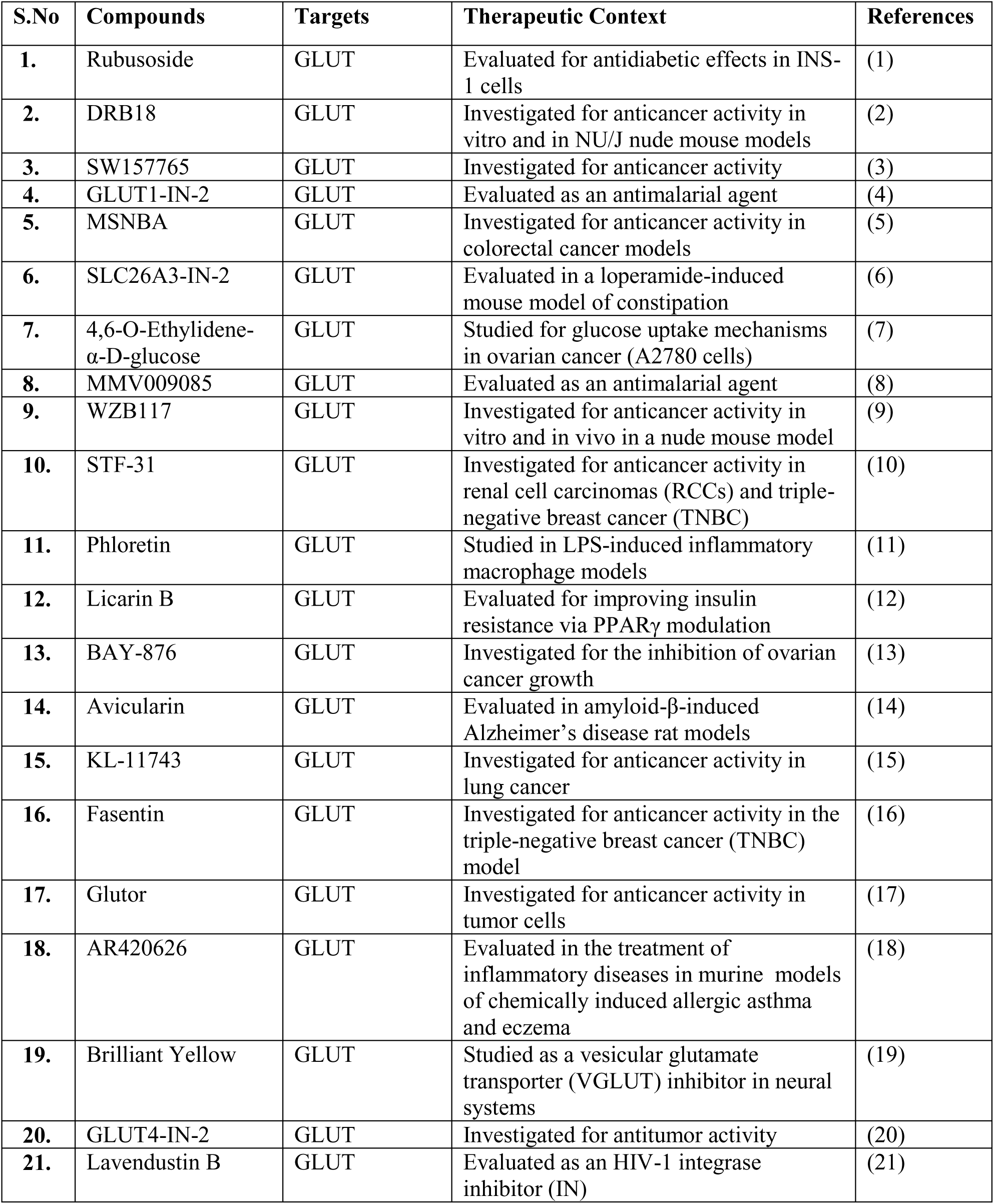
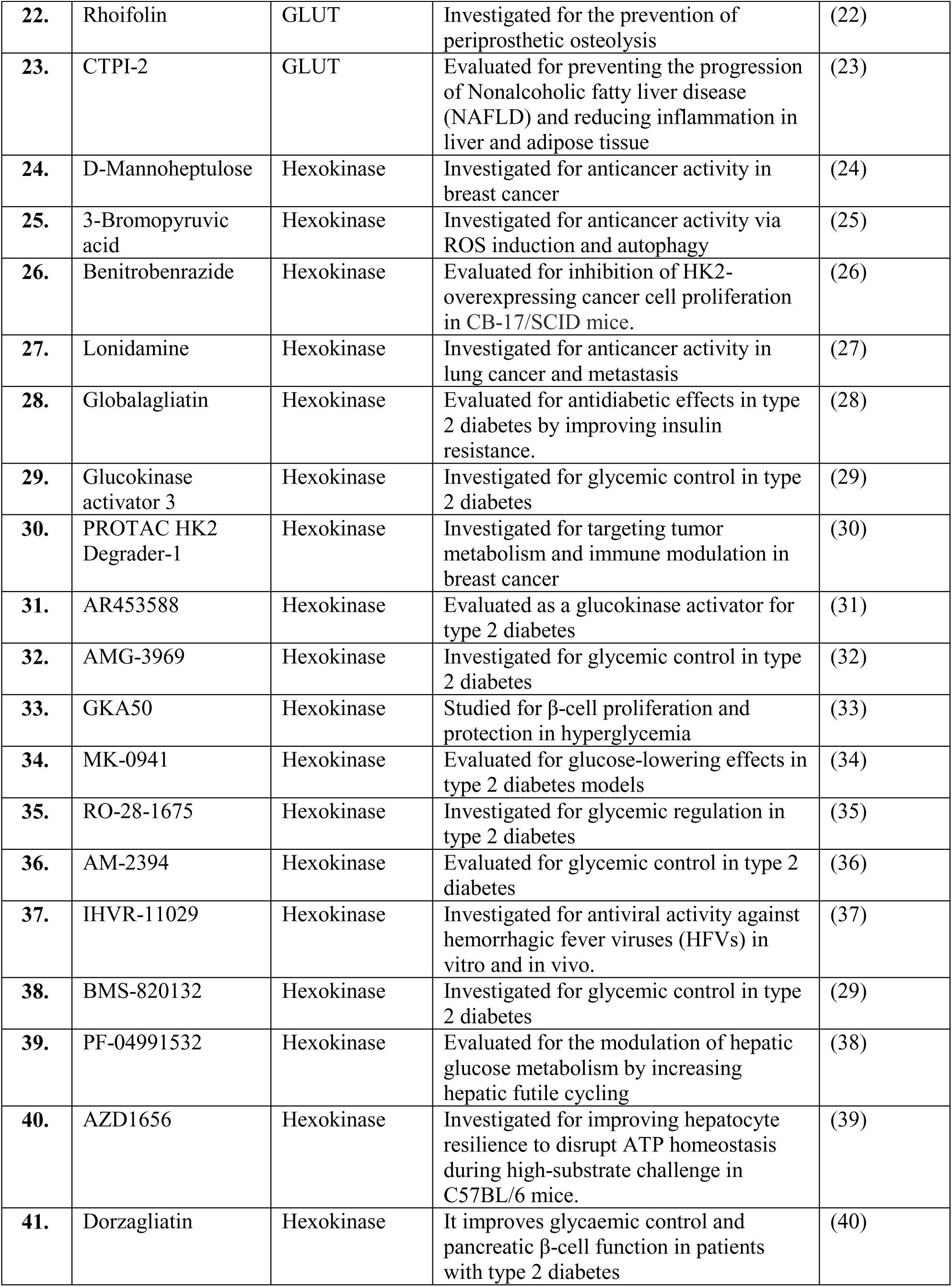
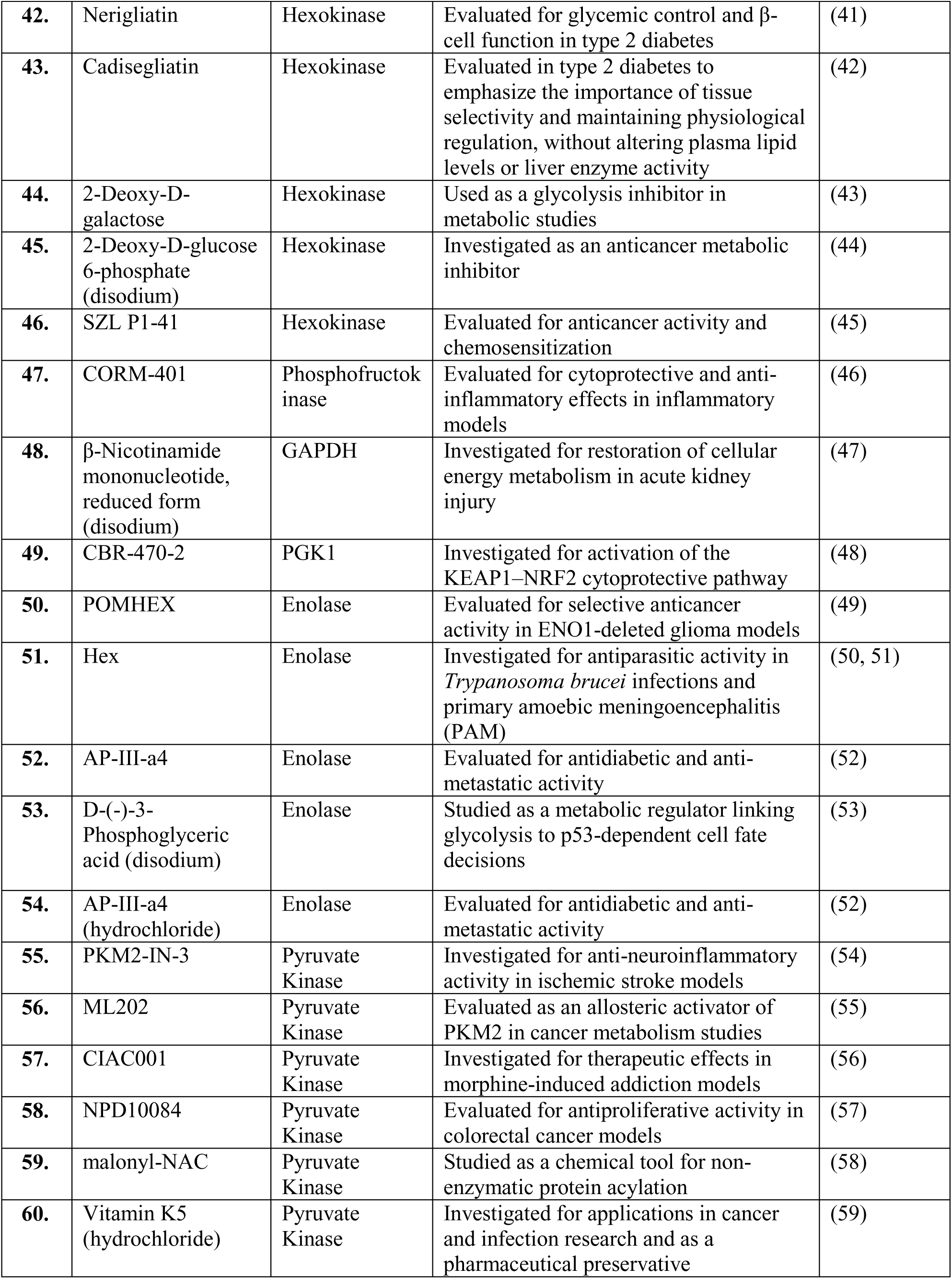
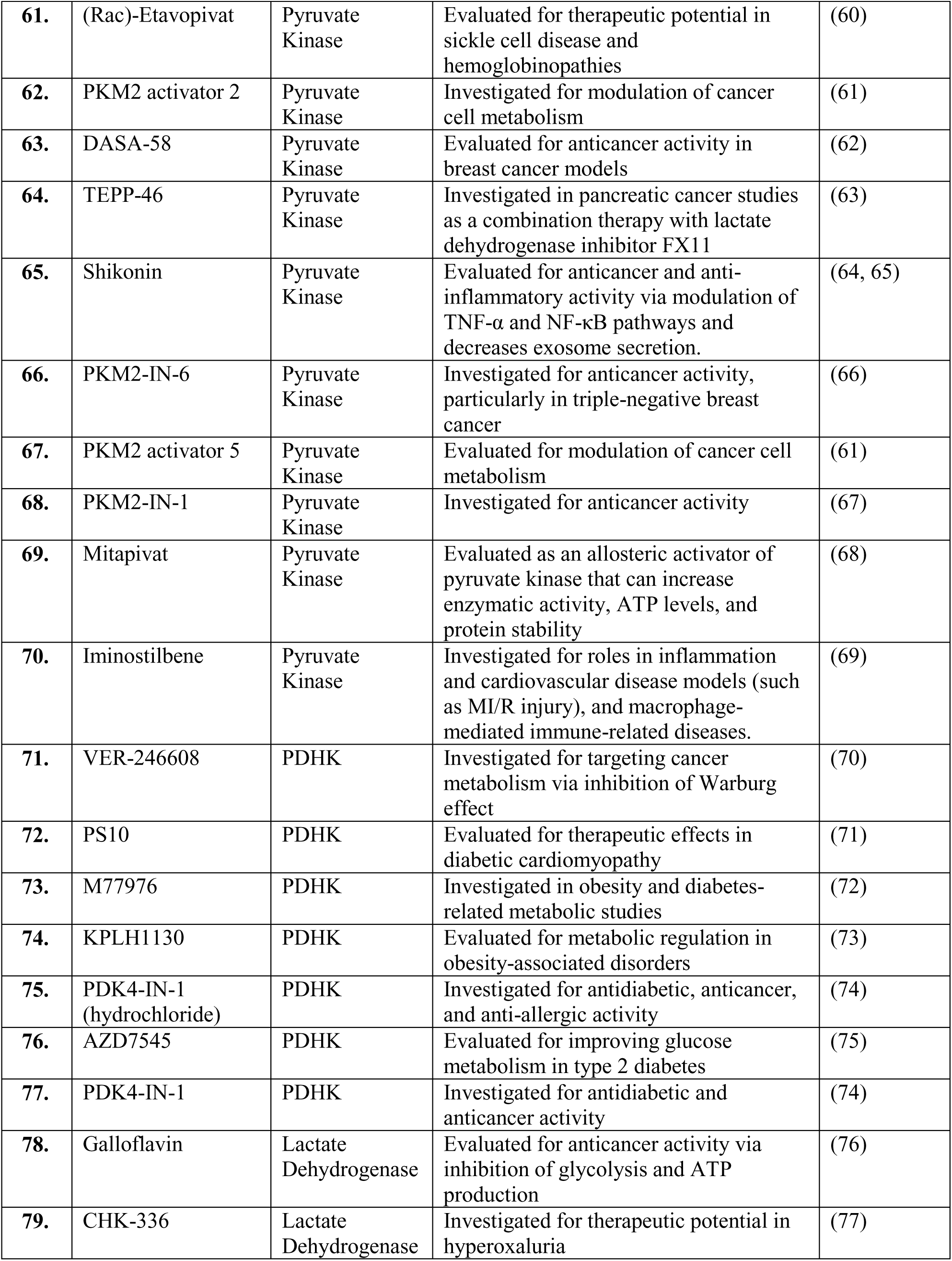
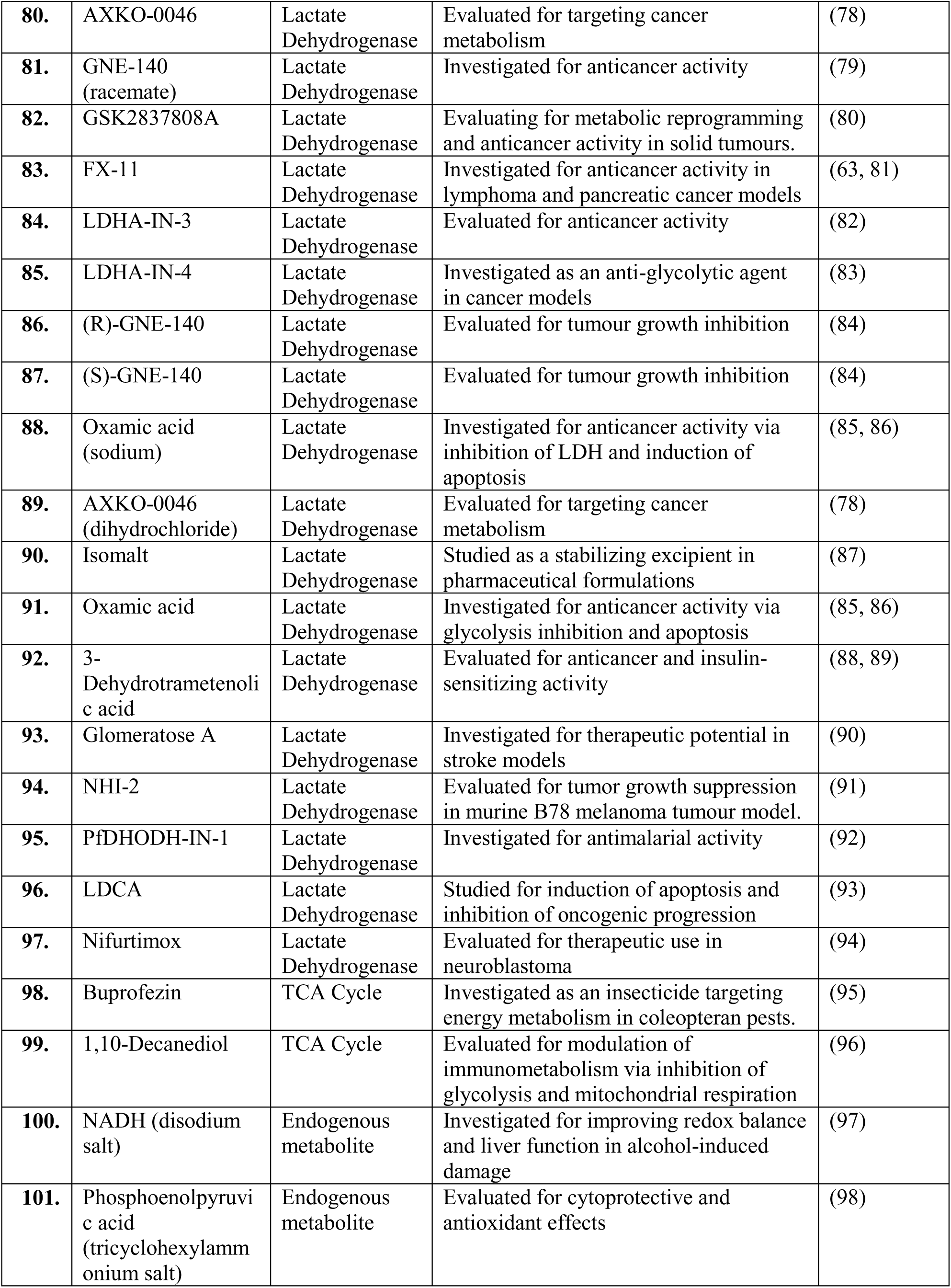
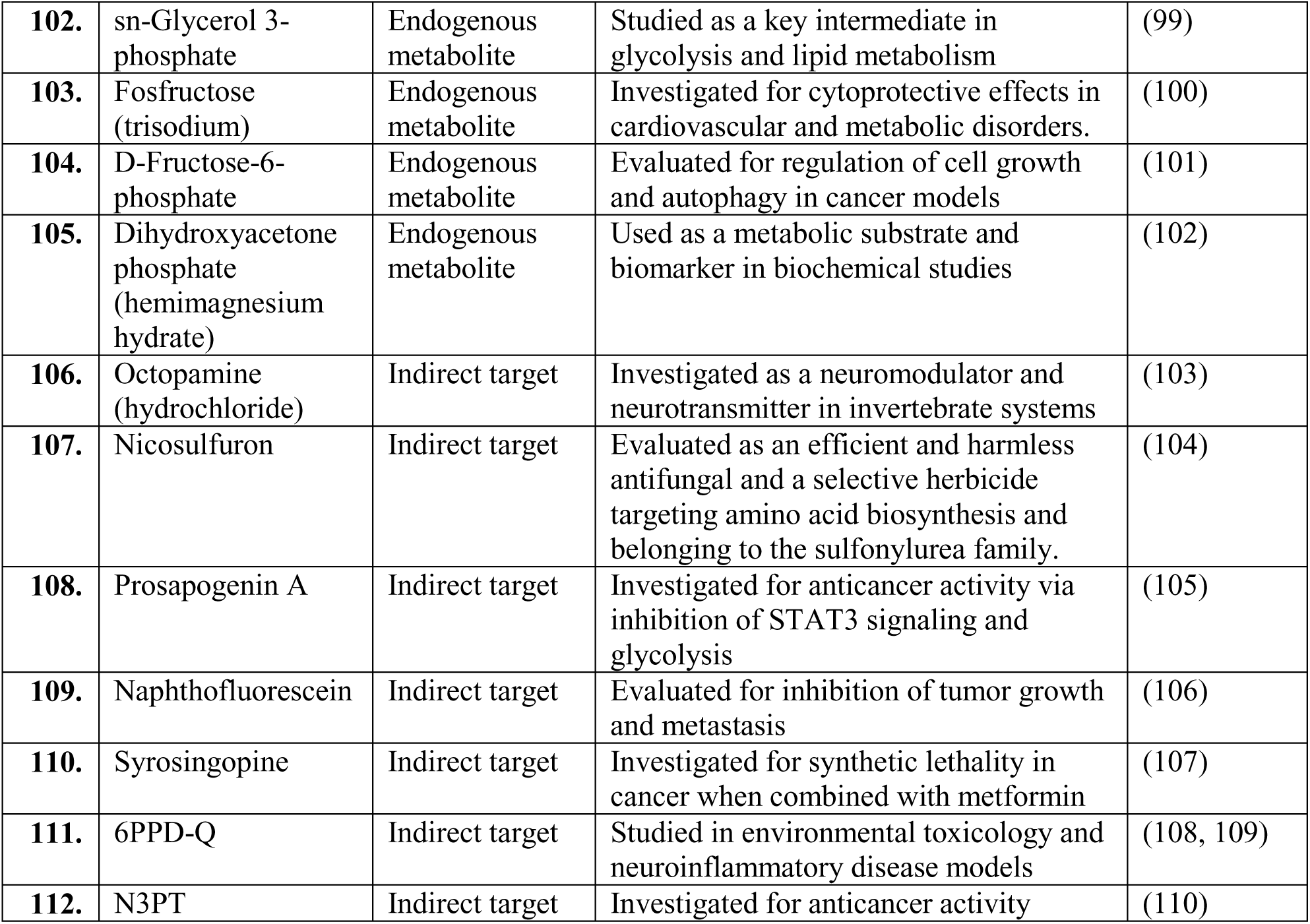
List of Glycolysis-targeting Compounds used in this Study.

